# Transferrin Receptor 1 is Required for CD4+ T Cell Iron-Dependent Response During Acute *Toxoplasma gondii* Infection

**DOI:** 10.1101/2023.06.28.546960

**Authors:** Stephen L. Denton, Tathagato Roy, Hunter K. Keplinger, Sai K. Ng, Jason P. Gigley

## Abstract

Elemental iron is an essential nutrient involved in many biological processes including infection and immunity. How iron impacts *Toxoplasma gondii* (*T. gondii*) *in vivo* and development of immunity during infection is unclear. We found that although iron is required for parasite proliferation *in vitro,* paradoxically, iron restriction *in vivo* increased parasite burdens during acute and persistent infection stages and decreased survival of mice. Iron restriction lowered IL-12 and IFN*γ* in spleen and serum, but ratios of myeloid cells and the number and function of Natural Killer cells were unchanged. Iron restriction significantly impaired the development of CD4+ and CD8+ T cell responses to *T. gondii* during replicating type II and attenuated vaccine strain *cps1-1* infection. Low iron conditions reduced the percent and absolute numbers of antigen experienced CD11a+CD49d+, functional IFN*γ*+, and CD62L-KLRG1+ effector T cells. Iron restriction also decreased vaccine efficacy of *cps1-1* strain against secondary lethal challenge. Antigen experienced CD4+ and CD8+ T cells both upregulated their iron transporter Transferrin receptor 1 (CD71) during infection regardless of iron restriction. Mice whose CD4+ T cells were deficient in CD71 had reduced CD4+ T cell antigen experience and polyfunctionality, yet CD8+ T cell responses remained intact, and their long term survival was not affected compared to wild type litter mate controls. This study highlights that iron acquisition by T cells is required for activation and vaccine induced long-term protection against *T. gondii*. Understanding how iron affects multiple immune compartments will be essential to define iron regulation of immunity to *T. gondii*.

## Introduction

*Toxoplasma gondii (T. gondii)* is an obligate intracellular protozoan parasite infecting about 30% of people world-wide which persists for life and poses a significant health risk (1–3). Understanding the factors that contribute to pathogenesis and development of immunity to this pathogen are critical to develop better vaccines and therapies to treat this infection. *T. gondii* has evolved to rely on scavenging nutrients from the host (4–6). Many of these nutrients are also required for proper development of immune responses to control this infection (7–9). The interplay between parasite nutritional scavenging and immune response development dependent on the same nutrients is poorly understood.

Essential metal iron is a nutrient that is utilized both by *T. gondii* and the development of immunity to the parasite. *T. gondii* is highly sensitive to exogenous control of iron levels (10, 11). Treatment of infected cells with IFN*γ* impairing parasite replication is rescued by iron salt and iron-bearing transferrin (12). A natural function of IFN*γ* treatment of cells is downregulation of iron uptake receptor Transferrin Receptor (CD71) (13); CD71 in turn is upregulated by *T. gondii* secretions (14). Modulating iron levels in mice infections has been shown to affect intestinal inflammatory damage as well as parasite load (15). However, the long term effects of iron modulation on development of protective immunity to *T. gondii* is not known.

Immunity to *T. gondii* is triggered by recognition of the parasite by TLR4 and 11 and the production of the proinflammatory cytokine IL-12 (16–20). IL-12 activates group 1 Innate Lymphoid Cells including Natural Killer (NK) Cells and ILC1 for the initial production of IFN*γ* required for early control of *T. gondii* (21–23). After presentation of *T. gondii* antigens to T cells by CD8a+ Dendritic cells, both CD4+ and CD8+ T cells mount a robust response producing high levels of IFN*γ* (24). CD8+ T cells, *via* their IFN*γ* production, emerge to become the dominant cell required for long-term protection against the infection. CD4+ T cells are required continuously to help maintain CD8+ T cell dependent protection (25, 26). Despite induction of a robust CD8+ T cell response, immunity to *T. gondii* is not sterilizing and the parasite escapes to develop a persistent infection in skeletal muscle and the central nervous system (27). Part of the inability of CD8+ T cells to clear the persistent infection is the development of immune exhaustion (27–29). The immune exhaustion of CD8+ T cells maybe partially dependent upon CD4+ T cells responding to the infection, highlighting that CD4+ T cells can be essential but also detrimental to long term CD8+ T cell dependent protection (26, 30). The immunological reasons why the parasite is not completely cleared is still not well defined. Available iron could be an important factor required in this process.

Iron-dependent roles in immunity include neutrophil extracellular trap formation (31), macrophage polarization (32, 33), dendritic cell maturation (34), and NK cell target recognition (35, 36). T Cell proliferation and IL-2R upregulation during activation is also dependent on intracellular iron levels (37) and CD71-dependent iron uptake (38). T-helper type 1 (Th1) but not Th2 cytokine production is also dependent on iron status (39, 40), indicating that relative iron levels could be a sensing mechanism for T-helper type differentiation. Much of what is known about iron and immunity is in the absence of infection. How iron regulates infection pathogenesis and development of immunity during infection including *T. gondii* infection *in vivo* is unclear.

We investigated how limiting iron during infection in a murine model impacted infection pathogenesis and immunity to *T. gondii*. Similar to previously published studies, we observed that limiting iron *in vitro* slowed the proliferation of the parasite. However, unlike other studies we observed that there was increased morbidity and mortality of the animals in iron limiting conditions, and the parasite disseminated better to the CNS to develop a much higher persistent cyst burden than normal iron conditions. To examine the role of iron in *T. gondii* resistance, we observed that splenic and serum production of IL-12p40 and IFN*γ* by 5 days after infection were inhibited by iron restriction, but the proportions of spleen leukocytes and NK production of IFN*γ* were unaffected. However, the expansion of antigen experienced CD4+ and CD8+ T cells responding to infection was dependent on iron availability. Consequently, the Th1 polyfunction (TNF*α* and IFN*γ* production) of the CD4+ T cells was inhibited by iron restriction in *T. gondii* infection. Inhibition of T cell responses by iron restriction was independent from antigen load and impacted development of long-term vaccine induced protection. CD4+ T cell specific expression of CD71 was intrinsically required for antigen experience and polyfunction of CD4+ and not CD8+ T cells, yet conditional CD71 knockout mice were not more susceptible to *T. gondii* infection. These results demonstrate the importance of iron in global development of protective immunity to *T. gondii* but also that iron-sensing by CD71 regulates CD4+ T cell activation is a critical component of long-term immunity to *T. gondii* infection.

## Materials and Methods

### Ethics Statement

All animal husbandry and experiments were performed by the Gigley Laboratory at the University of Wyoming in strict accordance to the guidelines approved by the University of Wyoming Institutional Animal Care and Use Committee.

### Mouse strains, parasite propagation, and infections

C57BL/6J (#000664), RAG1-/- (B6.129S7-Rag1^tm1Mom/J^, #002216), and CD4- CreER^T2^ (B6(129X1)-Tg(Cd4-cre/ERT2)11Gnri/J, #022356) mice were purchased from Jackson Laboratories. CD71^Flx/Flx^ mice were a kind gift from Mark Andrews, Harvard University. CD4-CreER^T2^ mice were crossed with CD71^Flx/Flx^ for two generations and the resulting progeny were interbred to produce CD4-CreER^T2^ +/-::CD71^Flx/Flx^. The genomes of these mice contained 75% C57BL/6 heritage from the CD4-CreER^T2^ parentage and 25% 129 heritage from the CD71^Flx/Flx^ parentage, thus, all experiments using these mice used internal littermate controls. Mouse genotypes were assessed by PCR from genomic DNA isolated from ear clips *via* digestion in DirectPCR Digestion Buffer + Proteinase K (Viagen Biotech) using the following primer pairs: CD4-CreER^T2^ Forward (0.5µM): 5’- GCG GTC TGG CAG TAA AAA CTA TC -3’, CD4-CreER^T2^ Reverse (0.5µM): 5’- GTG AAA CAG CAT TGC TGT CAC TT -3’, and separately TfR^Flx^ Forward (0.3µM): (5’-TTC AGT TCC CAG TGA CCA CA -3’), TfR^Flx^ Reverse (0.3µM): 5’- TCC TTT CTG TGC CCA GTT CT -3’. The CD4-CreER^T2^ transgene was detected by a 100nt PCR product. The 310nt lox-containing CD71 allele was differentiated from the 165nt wild-type CD71 allele by gel electrophoresis. Female mice aged 8-12 weeks old were used for all experiments.

Type II *T. gondii* ME49 strain was maintained by serial passage in CBA mice (Jackson Laboratories #000656). Cyst numbers were counted in 10µL of brain homogenate from previously infected animals placed on a glass microscope slide. Mice were inoculated intragastrically (i.g.) with 7-10 cysts in 200µL Phosphate-Buffered Saline (PBS) via oral gavage.

Type I *T. gondii* RH strain was maintained by routine passage in MRC5 fibroblasts in 1 % Dulbecco’s Modified Eagles Medium (1% DMEM) (1% Fetal Bovine Serum, 1X Penicillin/Streptomycin, 1X Amphotericin B). Upon lysis of MRC5 monolayers, tachyzoites were purified by 3µm filtration and enumerated on a hemocytometer. 1 X 10^3^ parasites were injected intraperitoneally (i.p.) in 200µL PBS. Attenuated Type I *T. gondii* strain *cps1-1* (CPS) was maintained by routine passage in MRC5 fibroblasts in 1% DMEM supplemented with 300mM uracil. Following filtration and enumeration similar to RH cultures, CPS was injected i.p. with a 27G needle at parasite concentrations as indicated in 1 X PBS.

### Drug Formulation and Treatment

Deferiprone (3-Hydroxy-1,2-dimethyl-4(1H)-pyridone “DFP”, Sigma-Aldrich #379409, CAS #30652-11-0) was dissolved at 10mg/mL in distilled water at 37°C and sterilized by filtration through a 0.2µm membrane. Each preparation was tested for Iron-Binding Units using a colorimetric Iron Assay Kit (Abcam Cat #ab83366) according to the manufacturer’s instructions. Briefly, a standard curve was created using a known amount of iron was prepared in Assay Buffer, and variable volumes of DFP preparations were added to replicates of the top standard. Iron reducer was added to every sample to measure total free iron in the solution. After addition of the iron probe, the absorbance of the samples was measured at 593nm and the amount of free iron remaining in the sample was extrapolated from the standard curve. The volume of DFP from each preparation required to reduce the iron in the samples by 50% was deemed an Iron-Binding Unit (U). DFP was administered at 75U per mouse intraperitoneally with a 27G needle every other day starting the day preceding infection, unless otherwise indicated.

Tamoxifen (Sigma-Aldrich #T5648, CAS #10540-29-1) was dissolved at 20mg/mL in corn oil (Sigma-Aldrich #C8267, CAS #8001-30-7) overnight by agitation at 37°C in the dark. Tamoxifen was administered at 100mg/kg to mice intraperitoneally with a 27G needle daily for the five days preceding infection.

### ELISA

Blood samples were collected by retroorbital bleeding of isoflurane anesthetized mice, followed by a coagulation period for 30 minutes at room temperature. Serum was collected after centrifugation. Splenic samples were suspended at 5*10^6^ cells per well in FACS Media and incubated for 48 hours at 37°C and 5% CO_2_. Cells were lysed by 3X freeze/thaw cycles at -80°C, and supernatant isolated for ELISA. Both serum and spleen samples were diluted at a volume of 1:20 for IL-12p40 and 1:15 for IFN*γ* ELISAs, which were performed according to manufacturer instructions using kits for IL-12p40 (Biolegend 431604) and IFN*γ* (Biolegend 430804).

### Parasite Burdens by Genome Equivalence

Genomic DNA from specified organs was purified from the samples using the DNeasy Blood & Tissue Kit (Qiagen 69506) according to manufacturer instructions. A standard curve of known quantity of RH gDNA was assessed alongside 100ng-400ng of sample DNA as template to query for the *T. gondii* unique gene B1 using SSO Advanced SYBR Master Mix (Bio-Rad 1725017) and 0.5µM primers (F: 5’- CGT CCG TCG TAA TAT CAG -3’, R: 5’- GAC TTC ATG GGA CGA TAT G -3’). The thermocycler conditions were 1: 95°C for 10 minutes, 2: 95°C for 5 seconds, 3: 60°C for 5 seconds, 4: 72°C for 5 seconds + plate read, 5: Cycle 2-4, 44 repetitions, 6: Melt Curve 65°C to 95°C with 0.5°C, 5 second increments + plate read. Parasite concentrations were extrapolated from linear regression of the standard curve.

### Plaque Assay

MRC5 fibroblasts were seeded into 6-well tissue culture plates and treated with DFP or ferric citrate as indicated immediately prior to infection. 100 RH parasites were inoculated per well and incubated for 7 days at 37°C and 5% CO_2_. Monolayers were fixed with 100% EtOH and stained with Plaque Assay Stain (125mL of 10% Crystal Violet in EtOH w/v + 500mL 1% Ammonium Oxalate w/v). Plaques were visualized by projection using a Bellco Glass, Inc. Slide Magnifier and manually counted. Plaque size was estimated as a measurement of A = *π*\*m((LA+TA)/4)^2^, where A = Area (mm2), m = magnification factor, LA = length of longest axis of plaque (mm), and TA = length of transverse axis perpendicular to the longest axis (mm). Representative images were taken with an Apple® iPhone Xr and background digitally removed with Adobe® Photoshop.

### Single Cell Suspension

For peritoneal exudate cells (PEC), the peritoneum of mice was exposed intact, and injected with 10mL Stain/Wash Buffer with EDTA (SWBE, 5% w/v Bovine Calf Serum and 2mM EDTA in PBS) *via* a 21G needle. Cells were dislodged by external agitation of the cavity during gradual injection of the SWBE and redrawn in the syringe. Spleens were extracted and crushed through a 70µm cell strainer with SWBE washing, followed by osmotic lysis of red blood cells (RBC) by suspension in 1mL 1X RBC Lysis Buffer (Tonbo Biosciences #TNB-4300-L100) for 3 minutes. RBC lysis was quenched by addition of 20mL SWBE, and samples were passed through a 70µm cell strainer. Single cell suspension concentrations were estimated with a hemocytometer.

### Flow Cytometry- Minimal Information About T Cell / Natural Killer Cell Assays (MIATA-MIANKA)

Single cell suspensions were aliquoted to a clear FACS V-bottom plate at one million cells per well. Cell viability was estimated using LIVE/DEAD Fixable Aqua dye (ThermoFisher L34957), followed by extracellular antibody staining in SWBE, fixation and permeabilization by BD Cytofix/Cytoperm (BD Biosciences 554714), and intracellular / secondary staining in BD Perm/Wash buffer, all according to manufacturer instructions. Samples were processed in Guava EasyCyte 12HT Flow Cytometer and analyzed in FlowJo v.9.7.7 software.

For NK Cell assays, wells were coated with 0.5µg per well of NK1.1 (PK136, BioXCell BE0036) overnight at 4°C the day prior to staining. After plating in coated wells, cells were stimulated at 37°C and 5% CO_2_ four 4 hours with 0.66X Protein Transport Inhibitor Cocktail (PTIC, eBioscience 00-4980-03) in FACS Media (Isocove’s DMEM with L-Glutamine and 25mM HEPES supplemented with 1X Penicillin/ Streptomycin, Amphotericin B, Non-Essential Amino Acids, 0.1mM Sodium Pyruvate, 0.2mM Glutamine XL, 10% Fetal Bovine Serum, and 0.1mM 2-mercaptoethanol). Extracellular stain included CD3-FITC (17A2, Biolegend 100204), CD335/NKp46-APC (29A1.4, Biolegend 137608), and CD49b-Biotin (DX5, Biolegend 108904). Intracellular / secondary stain included IFN*γ*-PE (XMG1.2, Biolegend 505806) and Streptavidin-PE/Cy7 (Biolegend 405206). cNK cells were boolean gated on live cells by Live/Dead Aqua, lymphocytes by scatter profile, and as CD3-, CD49b+, and NKp46+.

For T Cell phenotype assays, cells were plated and proceeded with direct *ex vivo* staining. Extracellular stain included CD4-BV421 (GK1.5, Biolegend 100438), CD8-PE/Cy5 (53-6.7, Biolegend 100710), CD11a-PE/Cy7 (I21/7, Biolegend 153108), CD62L-APC/Cy7 (MEL-14, Biolegend 104428), CD49d-PE (9C10/MFR4.B, Biolegend 103706), KLRG1-PE (2F1/KLRG1, Biolegend 138408), KLRG1-APC (2F1/KLRG1, Biolegend 138412), CD71/TfR1-APC (RI7217, Biolegend 113820), and intracellular stain Ki67-PE (16A8, Biolegend 652404). Cells were boolean gated on live cell by Live/Dead Aqua, lymphocytes by scatter profiling, and either CD4+/CD8- or CD4-/CD8+.

For T cell function assays, cells were plated in FACS media supplemented with 4µg Toxoplasma Lysate Antigen (TLA) per test and stimulated for 8 hours at 37°C and 5% CO_2_. For the last 4 hours of stimulation, PTIC was added to each well at a final concentration of 1X. Extracellular staining included CD4-BV421 (GK1.5, Biolegend 100438) and CD8-PE/Cy5 (53-6.7, Biolegend 100710), and intracellular staining included IFN*γ*-PE (XMG1.2, Biolegend 505806), TNF*α*-APC (MP6-XT22, Invitrogen 17-7321-82), and GZB-FITC (QA16A02, Biolegend 372296). TLA was produced by sonication of purified RH strain *T. gondii* parasites in PBS, and the concentration was determined by Bradford Assay. Cells were boolean gated on live cell by Live/Dead Aqua, lymphocytes by scatter profiling, and either CD4+/CD8- or CD4-/CD8+.

For fluorescence activated cell sorting, 20 million cells per tube were assessed for cell viability by Live/Dead Aqua and stained without fixation with CD3-PE (17A2, Biolegend 100206) and CD4-FITC (GK1.5, Biolegend 100406). Samples were processed in a BD FACSMelody Cell Sorter before immediate RNA isolation. Sorting strategy is defined in Figure S4.

For splenic profiling screens, extracellular stain proceeded immediately *ex vivo* and the following antibodies were used: CD3-BV421 (17A2, Biolegend 100228), CD19-BV605 (6D5, Biolegend 115540), NK1.1-FITC (PK136, Biolegend 108706), CD11b-APC (M1/70 Biolegend 101212), Ly6C-PE (HK1.4, Biolegend 128008), Ly6G-PE/Cy7 (1A8, Biolegend 127618), CD11c-FITC (N418, Biolegend 117306), and I-E/I-A-PE (M5/114.15.2, Biolegend 107608). Gating strategy is defined in Figure 3A.

### RNA Isolation and qRT-PCR

Samples collected in FACS experiments were pelleted and resuspended in TRI Reagent (Sigma T9424) for RNA Isolation according to manufacturer instructions. RNA was further purified by RNA Cleanup Protocol of RNeasy Kit (Qiagen 74104). RNA samples were quantified by UV absorbance and cDNA synthesis was performed on normalized mass (50ng P1 samples, 20ng P2 samples) of RNA according to manufacturer instructions with Superscript IV Reverse Transcriptase (Invitrogen 18091050). Equal volumes of cDNA samples were used in qRT-PCR reactions using Sso Advanced Universal SYBR Master Mix (Bio-Rad 1725271) and validated primers (41, 42): Actin Forward (0.5µM) 5’- ACC CAC ACT GTG CCC ATC TA -3’, Actin Reverse (0.5µM) 5’- CAC GCT CGG TCA GGA TCT TC -3’, CD71 Forward (0.5µM) 5’- TCA AGC CAG ATC AGC AAT TCT C -3’, and CD71 Reverse (0.5µM) 5’- AGC CAG TTT CAT CTC CAC ATG - 3’. The PCR protocol was 1: 95°C for 3:00, 2: 95°C for 0:15, 60°C for 0:15, 72°C for 1:20 + Plate Read, 5: Repeat 2-5 44X, 6: Melt Curve 65.0°C to 95.0°C, 0.5°C increment for 0:05 + Plate Read. mRNA ratios were estimated using the ΔΔCt method.

### Statistical Analysis

Statistics and graph generation were completed using Graphpad Prism v.9 and Microsoft Excel v16.54 software. Experiments are pooled from at least two independent technical repeats containing 3-5 replicates per experiment. Statistics were performed between two groups by Student’s t-test or between multiple groups by One-Way ANOVA, or by Kaplan-Meier Survival analysis. Significance is indicated by *p<0.05, **p<0.01, ***p<0.005, and ****p<0.001. Figures were assembled in Adobe Illustrator 2021, references in Endnote X8.

## Results

### Iron is an essential nutrient for Toxoplasma gondii

To study the roles of iron in *Toxoplasma gondii* pathogenesis and immunity, it was necessary to induce a rapid depletion of iron at the onset of an infection. To this end, we selected use of the small molecule cell permeable iron chelator deferiprone (DFP) (43). Based on lot-to-lot differences of iron chelation ability, we developed experimental reproducibility by normalization of iron-binding efficiency in each formulation. In this assay, the redox ability of free iron reacts with Ferense S, a chromogen to produce a colorimetric readout linearly correlated with free iron concentration. We challenged a known amount of iron, 10nM, with a range of DFP amounts by batch. The required mass of each formulation to decrease the free iron to 5nM, or 50% binding, was deemed an Iron-Binding Unit (U) (Figure 1A).

**Figure 1.**
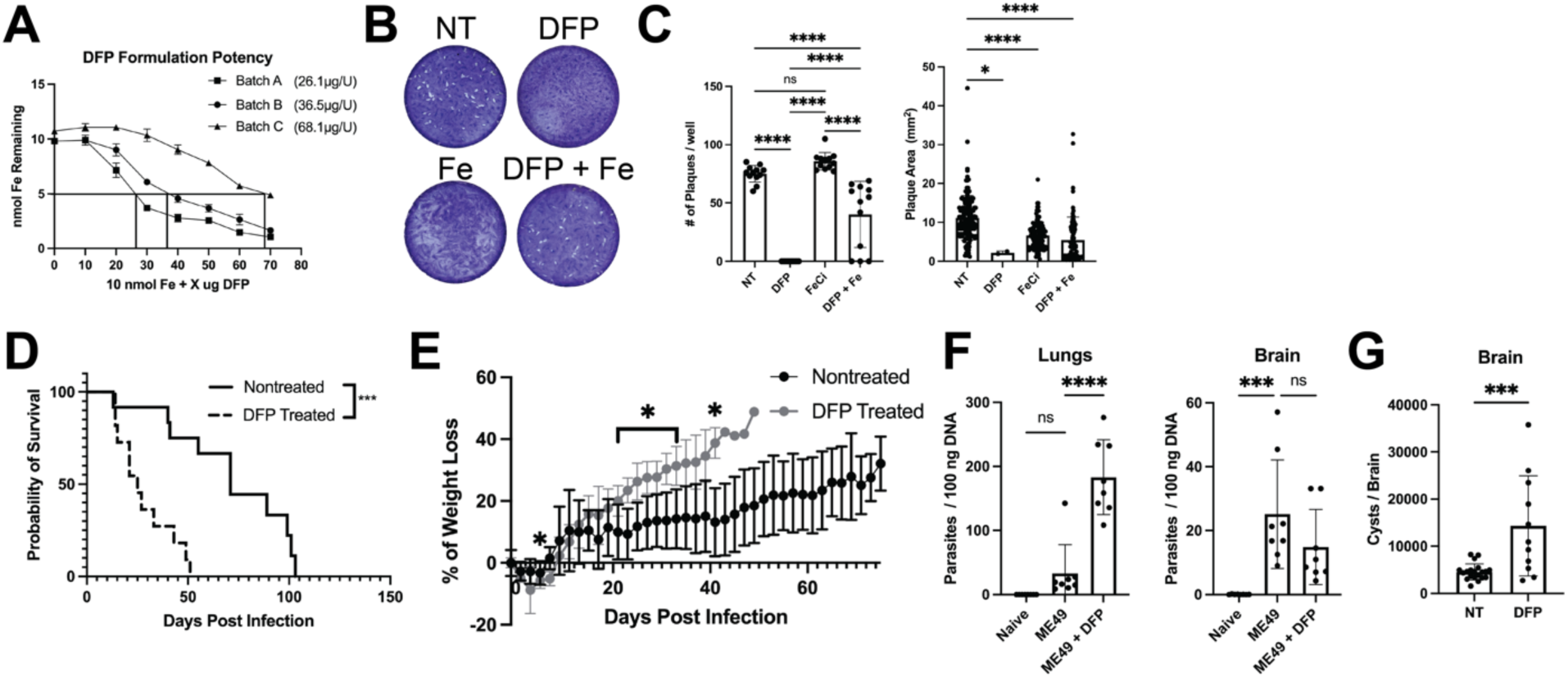
Iron restriction reduces *T. gondii* replication *in vitro* yet increases morbidity and mortality *in vivo*. **A)** Separate lots of DFP formulations were tested for Iron-Binding Units as described. Line graph shows titration curves of DFP used to determine 50% quenching of iron. **B)** Representative plaque assays are presented of RH strain *T. gondii* infection of fibroblasts that were non-treated (NT) or treated with 1.37U DFP, 30µg ferric citrate (Fe), or 1.37U DFP and 30µg ferric citrate (DFP + Fe). **C)** Bar graphs present pooled data from two independent replicates of N=6 each for total plaque number (left) and size of plaques (right, minimum 30 plaques measured per condition if possible). **D-H)** C57BL/6 mice infected with 8 cysts ME49 strain *T. gondii* i.g. and treated i.p. with 75U DFP every other day starting the day before infection were monitored or sacrificed at the timepoints indicated. **D)** Survival of infected mice after treatment with 75U DFP. Survival graph presents data pooled from three independent experiments of N=3-5 per group. **E)** Line graph presents percentage of weight loss over time of mice after treatment with 75U DFP. Data pooled from two independent repeats of N=4-5 per group. **F)** Parasite burden at 14DPI or time of death in infected organs after treatment with 75U DFP. Bar graphs present data from lung and brain pooled from two independent experiments of N=4 each. **G)** Brain cysts enumerated from infected mice treated with 75U DFP i.p. every other day starting on day -1 and surviving until day 35. Bar graph presents data pooled from four independent experiments but due to death of mice prior to day 35 the total sample size was 20 for NT and 11 for DFP treated groups. Bar graphs depict mean ± SD. Significance is denoted with * where p ≤ 0.05, ** where p ≤ 0.01, *** where p ≤ 0.005, **** where p ≤ 0.001, and NS where p > 0.05.

Dose titrating DFP into an infection setting showed a dose-dependent inhibition of parasite replication by plaque assay (Figure S1A) both in plaque number (Figure S1B) and plaque size (Figure S1C). The minimal dose tested that was sufficient to completely prevent plaque formation was 1.37U DFP, therefore, we tested this amount of DFP in a titration of a range of ferric citrate supplementation. A minimal dose of 20µg ferric citrate was sufficient to rescue the inhibition of parasite growth (Figure S1D). Lower doses of ferric citrate alone did not increase numbers of plaques observed (Figure S1E). However, at higher doses, which overcame DFP inhibitory effects, ferric citrate alone displayed toxicity to the host monolayer integrity (Figure S1A). This agrees with previous studies demonstrating the role of vacuolar Iron transporter (VIT) being important for regulating iron homeostasis in *T. gondii* (10). In summary experiments, DFP prevented *T. gondii* plaque formation in an iron-dependent manner (Figure 1B-C).

### Iron restriction “therapy” increases morbidity and mortality of T. gondii infection

Given the parasite-inhibitory properties of DFP *in vitro* we tested how iron chelation with DFP impacted infection pathogenesis *in vivo*. DFP is currently used in human patients as a safe iron overload therapy (43). During Type II ME49 strain infection in mice, treatment with DFP resulted in increased weight loss and mortality compared to non-treated controls (Figure 1D-E). The difference between iron chelation inhibition of proliferation *in vitro* and increased morbidity and mortality *in vivo* was surprising. Therefore we measured acute parasite burden in tissues of infected mice acute (14 DPI) and chronic (5 weeks PI, 5WPI) infection (20). DFP unexpectedly increased dissemination by 4-fold as assessed by parasite burden in the lung at 14DPI (Figure 1F), and by encystation in the brain at 5WPI (Figure 1G). These results indicate DFP restriction of iron impacts not only parasite proliferation, but dissemination and possibly development of immunity to *T. gondii*.

### IL-12 and IFNγ production is reliant on available iron in acute infection

IL-12 dependent IFN*γ* production is the key pathway for resistance to *T. gondii*, but also may contribute to inflammatory tissue damage (18, 22, 44). Therefore, changes in expression of these cytokines could correlate with the increased pathogenesis due to DFP-induced iron restriction. To determine how DFP treatment impacted development of immune control to *T. gondii* we compared IL-12 and IFN*γ* levels in naïve mice, ME49 infected mice and mice infected with ME49 and treated with DFP. As expected, *T. gondii* infection significantly increased IL-12 and IFN*γ* in spleen cells (IL-12: 17-fold, IFN*γ*: 32-fold) and serum (IL-12: 4-fold, IFN*γ*: 9-fold) compared to naïve animals (Figure 2). Iron restriction by DFP treatment significantly lowered the levels of these cytokines in both spleen cells and serum. These observations suggest that iron is a key factor in the activation of the immune response to *T. gondii*.

**Figure 2.**
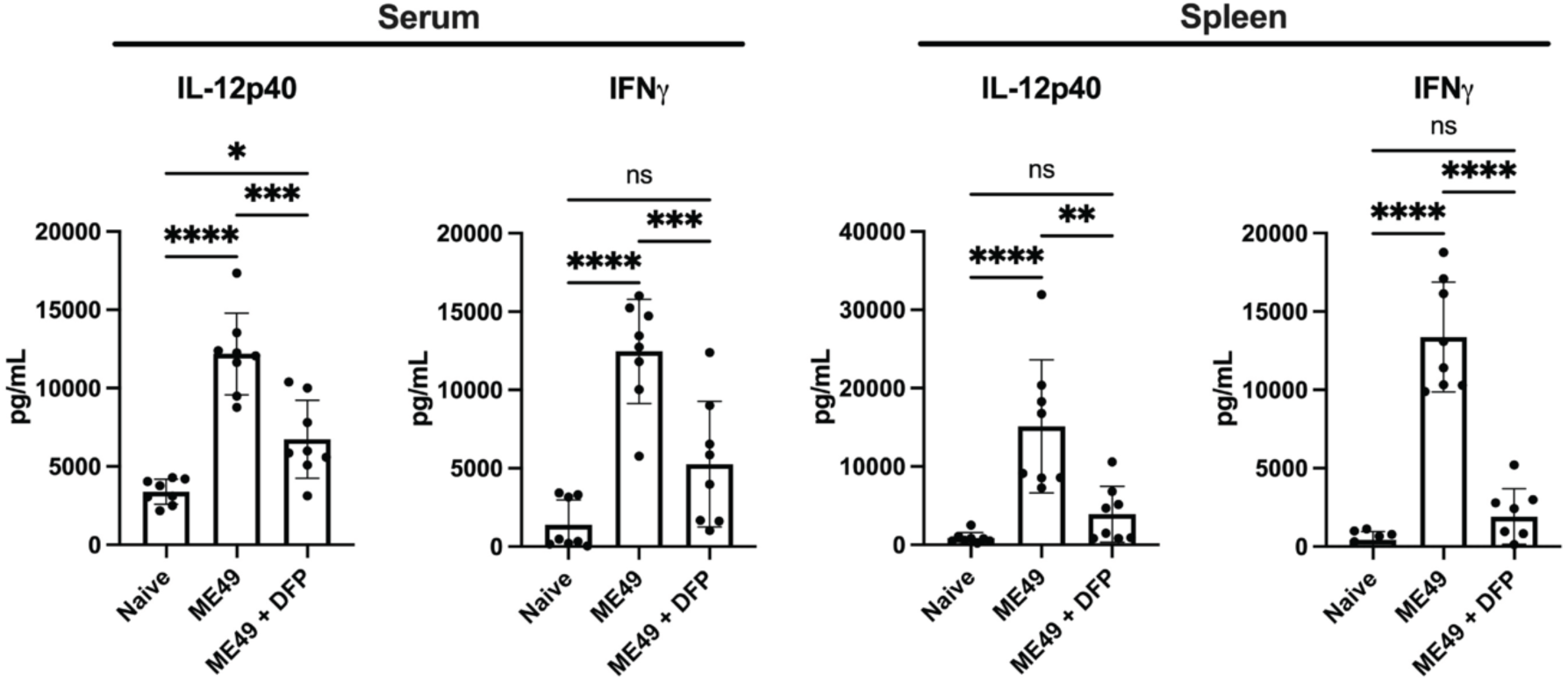
Iron restriction lowers IL-12 and IFNγ at 5DPI of *T. gondii* infection. Spleen cells and serum from C57BL/6 mice infected with 8 cysts ME49 I.G. and treated with 75U DFP were harvested at 5DPI. Bar graphs present results of ELISA measurements of serum levels and splenocyte production of IL-12p40 and IFN*γ* pooled from two independent experiments of N=4 each. All graphs depict mean ± SD. Significance is denoted with * where p ≤ 0.05, ** where p ≤ 0.01, *** where p ≤ 0.005, **** where p ≤ 0.001, and NS where p > 0.05

### Iron restriction does not impact splenocyte composition in acute infection

Because we observed significantly lower IL-12 and IFN*γ* during the early acute infection we further explored innate immune cell recruitment at this stage of infection. Neutrophils, Monocytes, Macrophages and Dendritic cells are all important producers of IL-12 (45, 46). Group 1 ILCs including natural killer cells and ILC1 are important early sources of IFN*γ* to initiate control of *T. gondii* (21–23). Analysis of spleen immune cell composition during acute ME49 infection with and without iron restriction by DFP revealed that restricting iron availability with DFP treatment (Figure 3A-B) did not have a significant impact on the frequencies of (CD19+ CD3-) B cells (Figure S2B), (CD19- CD3+ NK1.1-) T Cells (Figure S2C), (CD19- CD3- NK1.1+) NK Cells (Figure S2D), (Lineage- CD11b+ Ly6Chi Ly6G-) inflammatory monocytes (Figure S2E), (Lineage- CD11b+ Ly6Cint Ly6G+) neutrophils (Figure S2F), and (Lineage- CD11c+ MHCII+) Dendritic cells (Figure S2G). Notably, the total live lymphocyte numbers in spleen were significantly reduced at 5DPI only in iron restricted and infected animals, from 5.5-6*10^7^ cells to 4*10^7^ (Figure S2A). These results suggested that the overall composition of splenocytes could not explain the differences in morbidity, mortality and dissemination.

**Figure 3.**
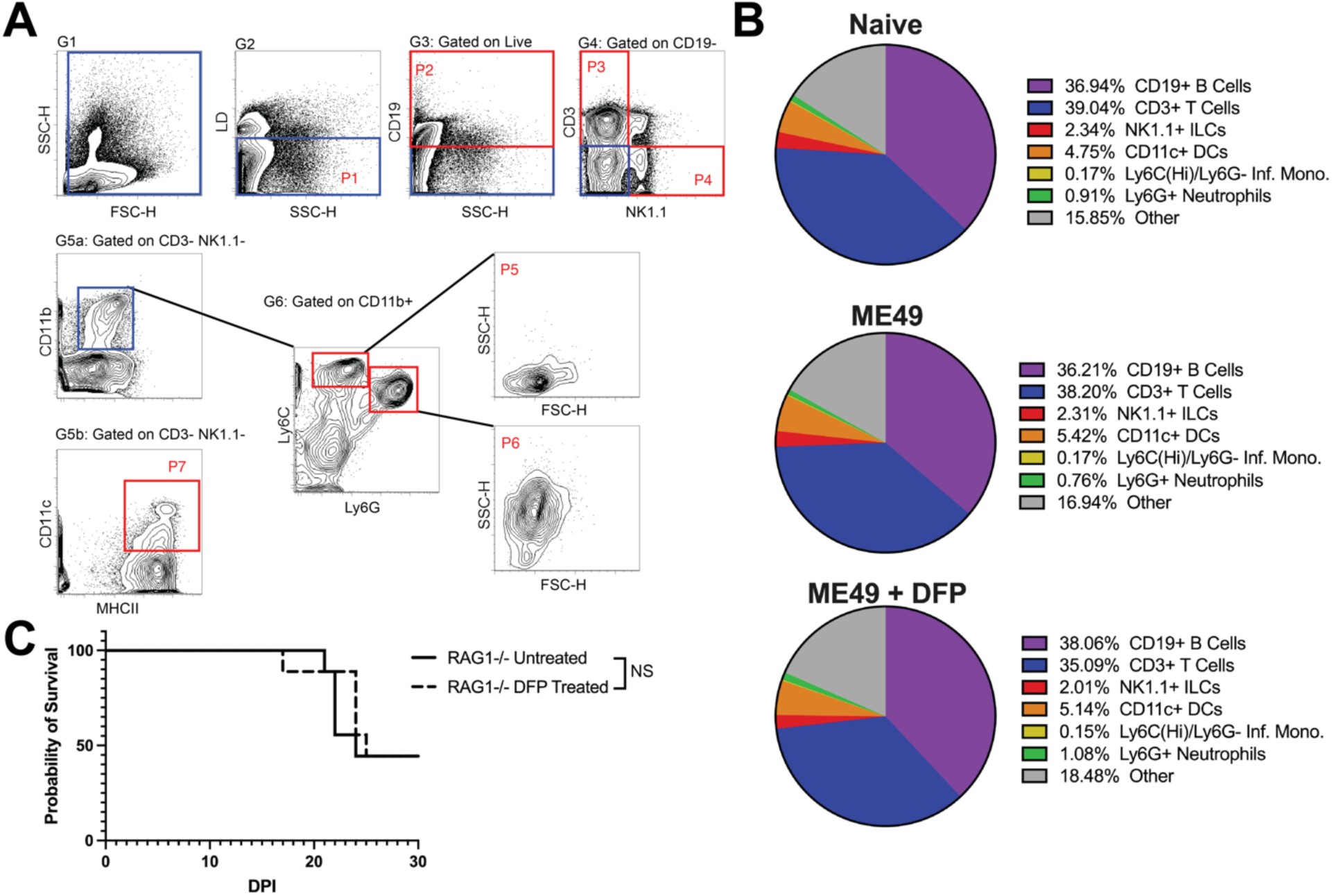
Iron restriction does not impact spleen cell proportions during early acute T. *gondii* infection. C57BL/6 mice were infected orally with 8 cysts ME49 and treated with 75U DFP i.p. on days -1, 1, and 3. Splenocytes were isolated on day 5 for flow cytometric analysis. **A)** Gating strategy presents Boolean gates (denoted G and in blue boxes) and population gates (denoted P and in red boxes). Lineage negative populations were gated as Live/Dead-, CD19-, CD3-, and NK1.1-. Population gates included P1 (Total Live Cells), P2 (Live, CD19+), P3 (Live, CD19+, CD3+, NK1.1-), P4 (Live, CD19-, CD3-, NK1.1+), P5 (Lineage-, CD11b+, Ly6CHi, Ly6G-, confirmed by side scatter profile), P6 (Lineage-, CD11b+ Ly6Cint, Ly6G+, confirmed by side scatter profile), P7 (Lineage-, CD11c+, I-A/I- E+). **B)** Pie charts depict the frequency means of indicated cell populations derived from raw data (Supplemental Figure 2) pooled from three independent experiments of N=4 per group. **C)** Survival of RAG1-/- mice infected orally with 8 cysts ME49 and treated with 75U DFP i.p. every other day starting the day before infection. Survival data is pooled from two independent experiments where NS indicates p > 0.05 by Kaplan-Meier analysis.

To further test if the innate cell composition is affected by iron restriction, we infected Rag1^tm1Mom^ (RAG1-/-), mice which are deficient in adaptive B or T cells (47), with cysts of ME49. RAG1-/- mice started to succumb to the infection starting at 2-3 WPI, however, iron restriction by DFP treatment during infection of these animals did not significantly change the mortality of the infection (Figure 3C). Taken together, these results suggest that the level of available iron impacts some innate immune cell function, but not cellular composition during acute *T. gondii* infection.

### Iron does not regulate NK cell number or IFNγ production

NK cells are critical for early T cell independent control of *T. gondii* infection (18, 21, 22, 48, 49). Even though there was no difference in mortality due to iron restriction in RAG1-/- animals who have an intact NK cell compartment, there was a significantly impaired systemic IFN*γ* production in WT animals. Therefore, we assayed NK cell (CD3- CD49b+ NKp46+) function in the presence or absence of iron restriction with DFP at 5 DPI in the spleen. The frequency and number of the NK cells dropped significantly during acute infection compared to naïve animals. However, the decrease in NK cells was similar regardless of DFP treatment (Figure 4A). NK cells were activated to produce IFN*γ* regardless of iron restriction by DFP suggesting that NK cells do not rely on available iron levels for their response (Figure 4B).

**Figure 4.**
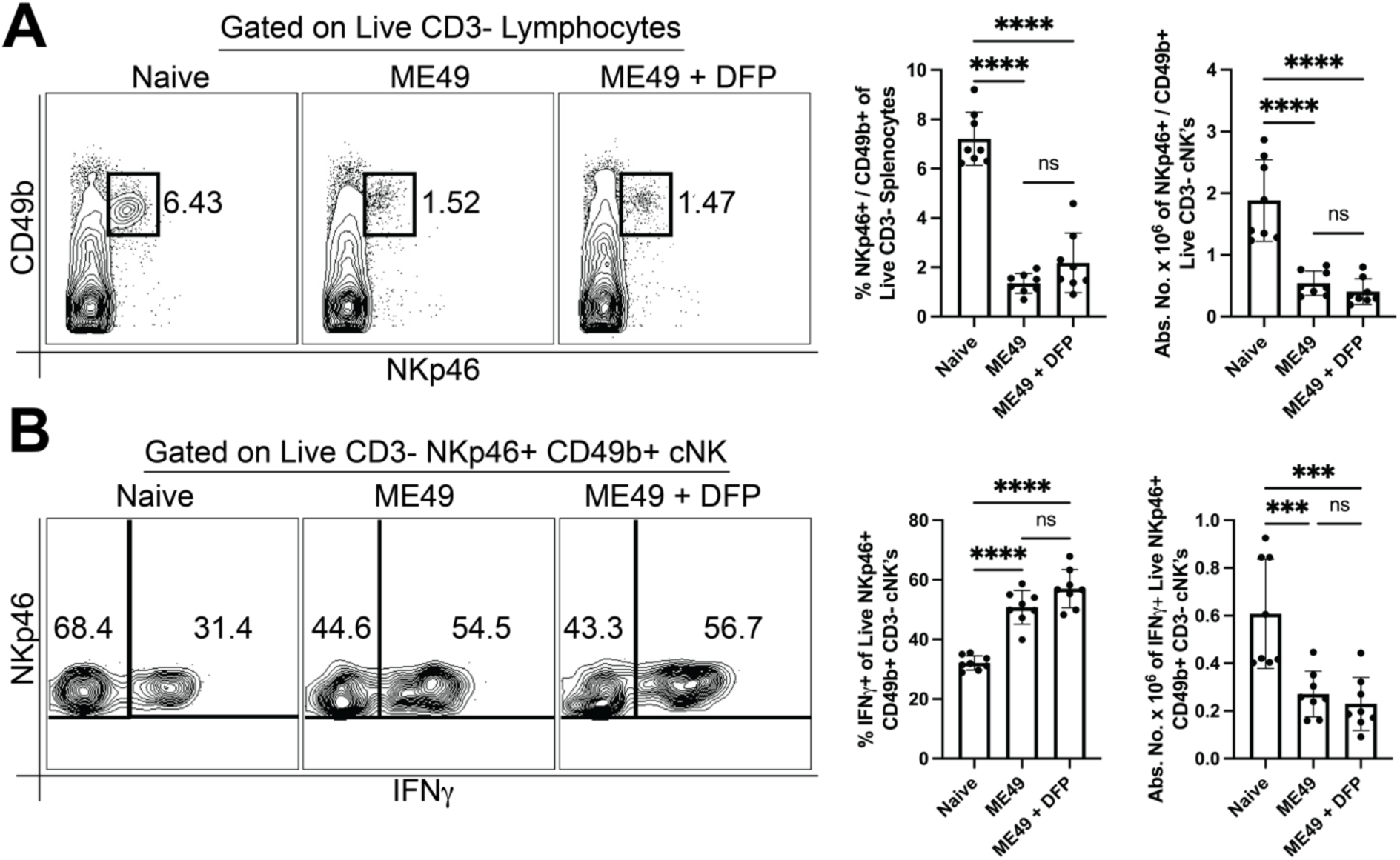
NK cell number and function are not affected by iron restriction during *T. gondii* infection. C57BL/6 mice were orally infected with 8 cysts ME49 and treated with 75U DFP i.p. on days -1, 1, 3. Splenocytes were stimulated with *α*-NK1.1 according to the MIANKA. **A)** Bar graphs and contour plots present frequency and absolute numbers of splenic cNK cells (Live, CD3-, CD49b+, NKp46+). **B)** Bar graphs and contour plots present frequency and absolute numbers of IFN*γ*-producing cNK cells (Live, CD3-, CD49b+, NKp46+, IFN*γ*+). Data pooled from two independent experiments of N=4 per group. All graphs depict mean ± SD. Significance is denoted with * where p ≤ 0.05, ** where p ≤ 0.01, *** where p ≤ 0.005, **** where p ≤ 0.001, and NS where p > 0.05.

### Iron is required to generate antigen experienced CD4+ and CD8+ T cells

Our results above suggest innate immune cell activity during iron restriction with DFP is relatively intact leading us to hypothesize that iron regulates and is essential for development of adaptive immune responses to *T. gondii*. We tested this hypothesis by examining the CD4+ and CD8+ T cell responses at the beginning of the adaptive response at 10 DPI with ME49 cysts by flow cytometry (50, 51). As expected, the frequency and number of total splenic helper CD4+ and cytotoxic CD8+ T cells was reduced after *T. gondii* infection compared to naïve animals (Figure 5A). In the iron-restricted infection group, this reduction in CD4+ and CD8+ T cells was comparable (Figure 5A). To better determine whether and how iron regulates T cell responses to *T. gondii* infection, antigen experienced T cells were assayed by flow cytometry. T cell upregulation of CD11a and tandem expression of CD49d is strongly correlated with markers of activation, thus, CD11a and CD49d (CD11a(Hi) CD49d+) serve as efficient surrogate markers of antigen experience (52). The frequency of the CD11a(Hi) CD49d+ population of antigen experienced CD4+ T cells increases from 5.7% in naïve animals to 47.3% in *T. gondii* infected animals without iron restriction (Figure 5B). Similarly, the frequency of CD11a(Hi) CD49d+ CD8+ antigen experienced T cells increased from 5.6% to 43.8% after *T. gondii* infection without iron restriction (Figure 5C). Absolute numbers of antigen experienced CD4+ and CD8+ T cells also increased significantly after infection in non-DFP treated animals compared to naïve controls (Figure 5B-C). However, the frequency of antigen experienced cells only reached 27.4% in CD4+ T cells and 22.1% in CD8+ T cells (Figure 5B-C). Moreover, the absolute number of antigen experienced CD4+ and CD8+ T cells under iron restriction was not significantly higher than in the naïve group. Iron restriction by treating animals with DFP during infection thus significantly inhibited increases in both the frequency and absolute number of antigen experienced CD11a(Hi) CD49d+ T cells compared to naïve and infected non-DFP treated animals.

**Figure 5.**
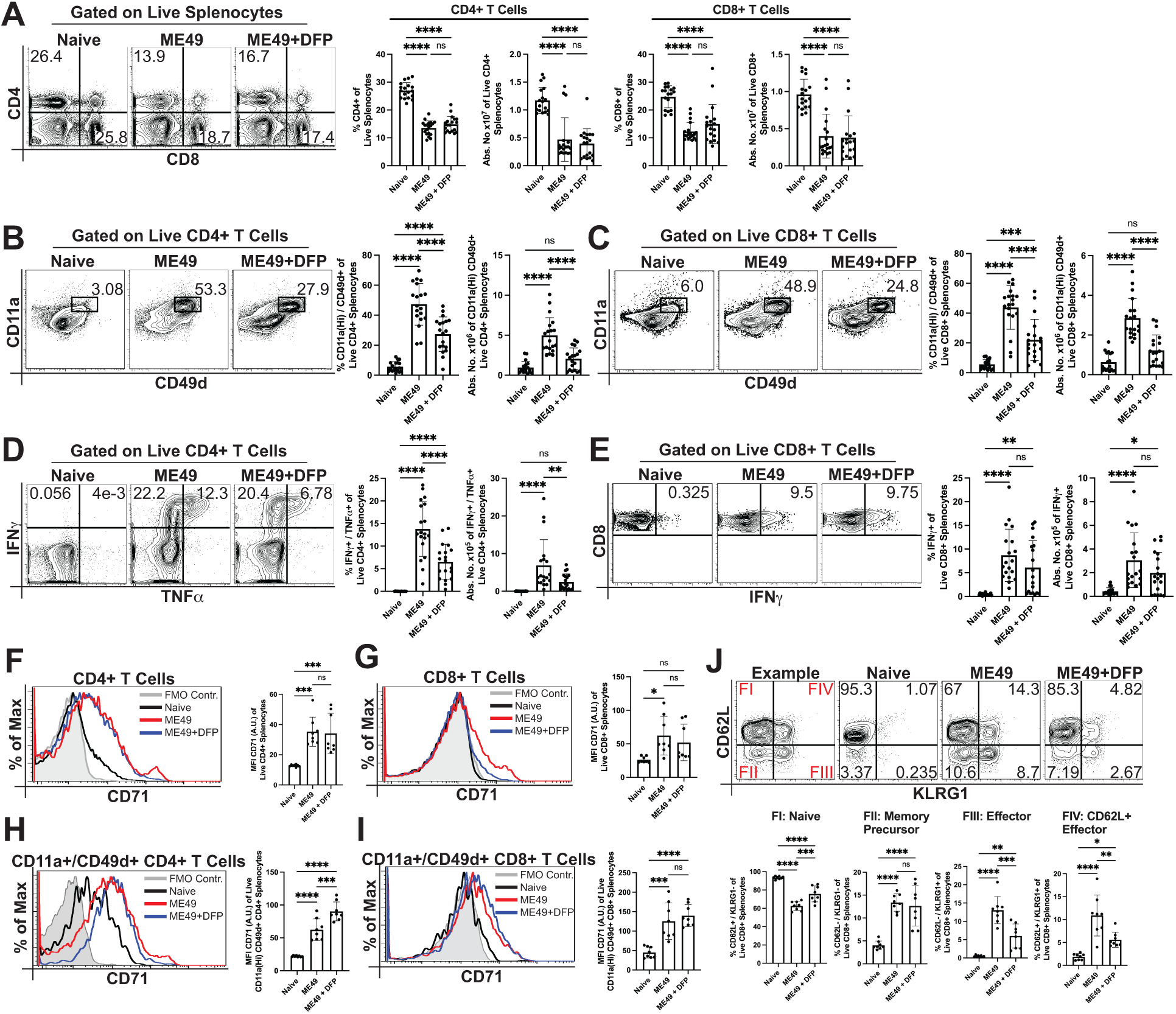
Iron is required for CD4+ and CD8+ T cell activation, expansion, function, and effector cell differentiation during Type II strain ME49 infection. C57BL/6 mice were orally infected with 8 cysts ME49 strain *T. gondii* and treated with 75U DFP i.p. every other day starting the day before infection. Splenocytes were prepared for stimulation with TLA or for direct *ex vivo* flow cytometry.. **A)** Direct *ex vivo* staining of CD4 and CD8 is represented by contour plots and bar graphs presenting frequency and absolute numbers of CD4+ T cells (Live/CD4+/CD8-) or CD8+ T Cells (Live/CD4-/CD8+). **B)** Direct *ex vivo* staining of CD11a and CD49d is represented by contour plots and bar graphs presenting frequency and absolute numbers of antigen experienced CD4+ T cells (Live/CD4+/CD8-/CD11aHi/CD49d+). **C)** Direct *ex vivo* staining of CD11a and CD49d is represented by contour plots and bar graphs presenting frequency and absolute numbers of antigen experienced CD8+ T cells (Live/CD4-/CD8+/CD11aHi/CD49d+). **D)** TLA-stimulated staining of IFN*γ* and TNF*α* is represented by contour plots and bar graphs of frequency and absolute numbers of polyfunctional CD4+ T Cells (Live/CD4+/CD8-/IFN*γ*+/TNF*α*+). **E)** TLA-stimulated staining of IFN*γ* is represented by contour plots and bar graphs of frequency and absolute numbers of functional CD8+ T Cells (Live/CD4- /CD8+/IFN*γ*+). **F-I)** Direct *ex vivo* staining of CD71 is represented by histograms and bar graphs presenting the geometric mean of CD71 intensity of F) Total CD4+ T cells (Live/CD4+/CD8-), G) Total CD8+ T cells (Live/CD4-/CD8+), H) Antigen experienced CD4+ T cells (Live/CD4+/CD8-/CD11aHi/CD49d+), and I) Antigen experienced CD8+ T cells (Live/CD4-/CD8+/CD11aHi/CD49d+). **J)** Direct *ex vivo* staining of KLRG1 and CD62L on CD8+ T cells is represented by contour plots and bar graphs presenting the frequency of FI: Naïve (CD62L+/KLRG1-), FII: Memory Precursor (CD62L-/KLRG1-), FIII: Short lived Effector (CD62L-/KLRG1+), and FIV: CD62L+ Effector (CD62L+/KLRG1+) subpopulations within the Live/CD4-/CD8+ population. All data pooled from 2-5 independent experiments of N=3-5 per group. All graphs depict mean ± SD. Significance is denoted with * where p ≤ 0.05, ** where p ≤ 0.01, *** where p ≤ 0.005, **** where p ≤ 0.001, and NS where p > 0.05.

### Iron is required for functional CD4+ T cell polyfunctional response

Due to the inhibitory effect of iron restriction on increases in T cell antigen experience, it was likely that the functions of these cells were also inhibited. Therefore we measured function of CD4+ (IFN*γ* and TNF*α*) and CD8+ T cells (IFN*γ*) *ex vivo* from mice infected with ME49 and treated or not with DFP to restrict iron (53–57). The frequency of polyfunctional IFN*γ*+ / TNF*α*+ CD4+ T cells increased from 0.017% to 13.8% after infection (Figure 5D). Additionally, the frequency of IFN*γ*-producing CD8+ T cells increased from 0.44% in naïve to 8.7% after infection (Figure 5E). The inhibitory effect of iron restriction on antigen experience in CD4+ T cells translated to inhibited function, as the frequency of polyfunctional CD4+ T cells was 6.5%, significantly lower than untreated infected animals, but still significantly higher than naïve animals (Figure 5D). In contrast, iron restricted animals still mounted a functional CD8+ T cell response, as the frequency of IFN*γ*+ CD8+ T cells (6.1%) was not significantly different than in untreated infection (Figure 5E).

### Antigen experienced CD4+ and CD8+ T cells upregulate CD71 expression

CD71 mediates iron uptake in most mammalian cell types, including immune cells (58). Additionally, CD71 is both upregulated and required for T cell activation *in vitro* (59–61). To link the inhibitory effect of iron restriction on T cell responses to T cell intrinsic iron uptake, we measured CD71 expression levels on CD4+ and CD8+ T cells during *T. gondii* infection using flow cytometry (Figure 5F-I). We observed a significant 2.7-fold increase in CD71 MFI on CD4+ T cells and a 2.4-fold increase on CD8+ T cells during *T. gondii* infection. In iron restricted animals, CD71 MFI of total CD4+ or CD8+ T cells was not significantly different compared to untreated infection groups (Figure 5F-G). We next measured CD71 MFI in antigen experienced CD11dHi CD49a+ T cells. Similar to total CD4+ and CD8+ T cells, CD71 increased in expression after *T. gondii* infection on CD11dHi CD49a+ antigen experienced CD4+ and CD8+ T cells compared to naïve conditions (Figure 5H-I). The MFI of CD71 was significantly higher on antigen experienced CD4+ and CD8+ T cells on a per cell basis (Figure 5H-I) than total CD4+ and CD8+ T cells (Figure 5F-G) from both non-DFP treated and iron restricted DFP treated infected mice. Iron restriction by DFP treatment also significantly increased CD71 MFI on antigen experienced CD4+ T cells compared to cells from non-treated infected animals (Figure 5H). Interestingly, iron restriction by DFP treatment did not impact the CD71 MFI of antigen experienced CD8+ T cells (Figure 5I). These results show that CD71 is upregulated on T cells during *T. gondii* infection, and that iron restriction by DFP treatment further upregulates CD71 only in antigen experienced CD4+ T cells.

### Iron restriction reduces frequency of effector CD8+ T cell subsets

Iron restriction significantly decreased the number of antigen experienced CD8+ T cells during *T. gondii* infection. To provide control of the parasite, CD8+ T cells differentiate from naïve (CD62L+KLRG1-, FI) into memory precursor (CD62L-KLRG1-, FII), short lived effector cell (CD62L-KLRG1Hi, FIII), and CD62L+ effector cells (CD62L+KLRG1+, FIV) (Figure 5J) (62). Therefore we measured CD8+ T cell differentiation in the presence or absence of iron restriction by DFP treatment during *T. gondii* infection defined by functional output (Figure 5J). The frequency of the naïve FI population decreased after infection from 93.55% of naïve animals to 62.65% in normal iron condition (non-DFP treated) animals. In the iron restricted (DFP-treated) group, there was a significantly greater proportion of CD8+ T cells remaining naïve at 75.73% compared to non-treated infected mice. After infection in non-DFP treated mice, CD8+ T cells differentiated into FII memory precursor subset (CD62L- KLRG1-, 13.38%), the FIII effector subset (CD62L- KLRG1+, 13.06%), and the FIV CD62L+ effector subset (CD62L+ KLRG1+, 10.89%). Iron restriction by DFP treatment inhibited differentiation of FIII (6.01%) and FIV (5.61%) effector subsets, both of which were significantly lower than in untreated infection and higher than naïve mice. However, the proportion CD8+ T cells belonging to the FII memory precursor subset (12.65%) was not significantly different between the DFP-treated or non-treated infection groups.

### Reduced T cell activation during iron restriction is independent of antigen load and occurs during vaccination

One of the reasons we may observe decreased antigen experienced CD4+ and CD8+ T cells and CD4+ T cell polyfunctional responses is the lack of sufficient *T. gondii* antigen due to the possibility that iron restriction slows *T. gondii* growth *in vivo*. The quantity and quality of antigen load is linked to differences in T cell activation and protection in *T. gondii* infection (25, 63, 64). Therefore to control for differences in antigen levels that might cause defects in T cell responses during DFP induced iron restriction, we immunized mice i.p. using 1 X 10^6^ tachyzoites of the non-replicating vaccine strain of *T. gondii,* CPS (24, 65) with or without DFP induced iron restriction. We analyzed the CD4+ and CD8+ T cell responses at the site of infection (PEC, Figure 6) and the systemic response on day 7 p.i. (spleen, Supplemental Figure 3).

**Figure 6.**
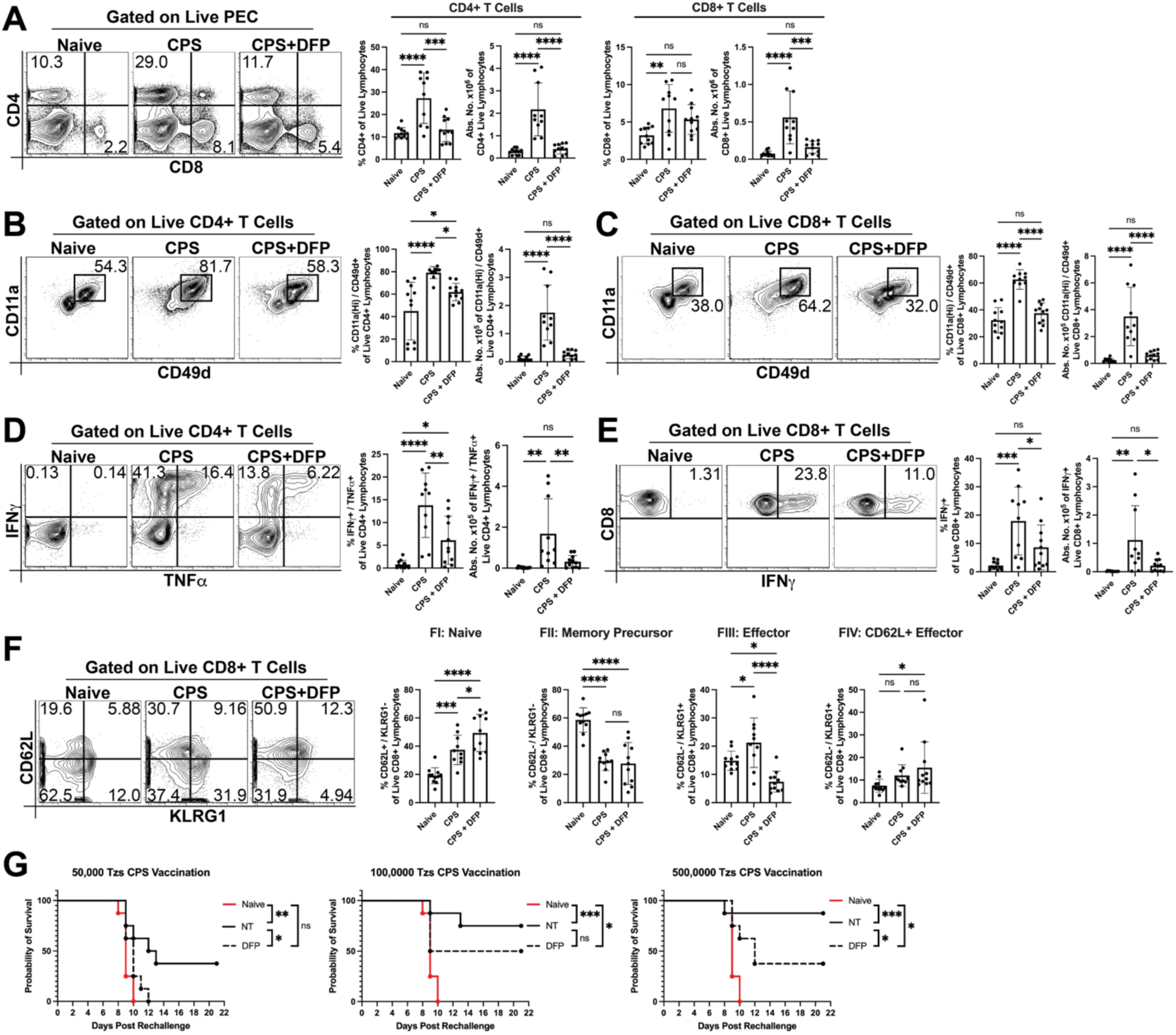
Inhibition of CD4+ and CD8+ T cell activation, expansion, function, and effector cell differentiation by iron restriction during vaccination is independent of antigen load. A-F) C57BL/6 mice were infected with 1 X 10^6^ tachyzoites CPS i.p. and treated with 75U DFP as described. At 7DPI, peritoneal Exudate Cells (PEC) were prepared for simulation or direct *ex vivo* staining. **A)** Direct *ex vivo* staining of CD4 and CD8 is represented by contour plots and bar graphs presenting frequency and absolute numbers of CD4+ T cells (Live/CD4+/CD8-) or CD8+ T Cells (Live/CD4-/CD8+). **B)** Direct *ex vivo* staining of CD11a and CD49d is represented by contour plots and bar graphs presenting frequency and absolute numbers of antigen experienced CD4+ T cells (Live/CD4+/CD8- /CD11aHi/CD49d+). **C)** Direct ex vivo staining of CD11a and CD49d is represented by contour plots and bar graphs presenting frequency and absolute numbers of antigen experienced CD8+ T cells (Live/CD4-/CD8+/CD11aHi/CD49d). **D)** TLA-stimulated staining of IFN*γ* and TNF*α* is represented by contour plots and bar graphs presenting frequency and absolute numbers of polyfunctional CD4+ T Cells (Live/CD4+/CD8-/IFN*γ*+/TNF*α*+). **E)** TLA-stimulated staining of IFN*γ* is represented by contour plots and graphs presenting frequency and absolute numbers of functional CD8+ T Cells (Live/CD4-/CD8+/IFN*γ*+). **F)** Direct *ex vivo* staining of KLRG1 and CD62L on CD8+ T cells is represented by contour plots and graphs presenting the frequency of FI: Naïve (CD62L+/KLRG1-), FII: Memory Precursor (CD62L-/KLRG1-), FIII: Short lived Effector (CD62L-/KLRG1+), and FIV: CD62L+ Effector (CD62L+/KLRG1+) subpopulations within the Live/CD4-/CD8+ population. **G)** C57BL/6 mice were injected with 1 X 10^4^ (left), 5 X 10^4^ (middle), or 1 X 10^5^ (right) tachyzoites of CPS strain *T. gondii* while being treated or not with 75U DFP for the first three weeks of vaccination. DFP treatment was then stopped for the last 2 weeks of vaccination allowing iron levels to recover. Graphs present survival of mice lethally challenged with 1 X 10^3^ RH strain *T. gondii* two weeks after cessation of DFP treatment. All flow cytometry data are pooled from three independent experiments of N=3-4 per group. All bar graphs depict mean ± SD. Vaccination studies were conducted twice. Significance is denoted with * where p ≤ 0.05, ** where p ≤ 0.01, *** where p ≤ 0.005, **** where p ≤ 0.001, and NS where p > 0.05.

In the PEC, the frequency and absolute number of total CD4+ and CD8+ T cells increased after vaccination (Figure 6A). Iron restriction using DFP significantly reduced those increases and were not significantly greater than naïve mice (Figure 6A). The frequency and number of antigen experienced (CD11aHi CD49d+) CD4+ and CD8+ T cells were then measured in the PEC of vaccinated mice with and without DFP iron restriction. In naïve animals, there was a high baseline frequency of CD11a(Hi) / CD49d+ antigen experienced CD4+ and CD8+ cells. The frequency and absolute number of antigen experienced CD4+ and CD8+ T cells significantly increased in response to vaccination with CPS compared to naïve controls (CD4+ frequency: 2.33-fold, number: 7.31-fold and CD8+ frequency: 2.13-fold, number: 7.43-fold). In contrast, DFP induced iron restriction resulted in a near complete inhibition of increased antigen experienced CD4+ T cell populations with only 1.13-fold (frequency) and 1.38-fold (number) non-significant increases after vaccination (Figure 6B). Similarly, DFP induced iron restriction inhibited CD8+ T cell populations with 1.67-fold (frequency) and 2.15-fold (number) non-significant changes compared to unvaccinated groups (Figure 6C). Consequently, we measured CD4+ T cell polyfunction (IFN*γ*+/TNF*α*+) and CD8+ T cell single function (IFN*γ*+) after CPS vaccination. Both the frequency and absolute number of single and polyfunctional CD4+ and CD8+ T cells were significantly increased after vaccination in the PEC. DFP induced iron restriction significantly reduced CD4+ and CD8+ T cell functions in both frequency and absolute number, and for CD8+ T cells, frequency and absolute number of functional cells were not significantly different from naïve animals (Figure 6D-E).

We next measured the ratio of naïve (CD62L+KLRG1-, FI), memory precursor (CD62L-KLRG1-, FII), short lived effector (CD62L-KLRG1+, FIII), and CD62L+effector (CD62L+KLRG1+, FIV) CD8+ T cells in PEC of naïve, CPS immunized/non-treated and CPS immunized/DFP-treated iron restricted mice (Figure 6F). We observed an increase in naïve CD8+ T cells (FI) after CPS vaccination compared to non-immunized animals and iron restriction further increased the frequency of these cells compared to naïve and CPS alone. Memory precursor CD8+ T cells (FII) decreased significantly after CPS vaccination with or without DFP treatment and DFP did not affect this decrease. The frequency of effector CD8+ T cells (FIII) increased significantly after CPS vaccination compared to naïve animals and DFP induced iron restriction significantly decreased this cell population to lower levels than naïve and CPS infected mice. There was only a moderate effect on CD8+ CD62L+ effector T cells with CPS immunization and CPS with DFP treatment compared to naïve mice.

In the spleen, the response to CPS is more subtle due to the localized immune response induced by this parasite strain in the PEC (62, 66). Measuring the CD4+ and CD8+ T cell responses in the periphery revealed that even though the frequency and absolute number of total T cells were unchanged (Figure S3A), antigen experience and function increased significantly after vaccination (Figure S3B-E). Similar to PEC these increases were inhibited in DFP induced iron restriction conditions and were not greater than naïve mouse cells. These results demonstrate that iron-restriction in CPS vaccination when antigen availability is normalized still significantly decreases T cell responses similar to ME49 infection, revealing the role of iron as a critical factor involved for the proper activation of T cell immunity to *T. gondii* independent of antigen level.

### Iron-restricted inhibition of adaptive immunity also reduces vaccine induced long-term protective immunity against lethal T. gondii rechallenge

Iron-restriction inhibits CD4+ and CD8+ T cells responses during primary infection of both ME49 and CPS strains. How iron impacts development of long term memory and control of *T. gondii after* vaccination with CPS has not been tested. CPS vaccination induces robust T cell memory that is highly protective against lethal challenge (25). Mice were immunized with a dose titration of CPS (5 X 10^4^, 1 X 10^5^, and 5 X 10^5^) tachyzoites and treated or not with DFP for the first three weeks after vaccination. Mice were then allowed to recover without DFP treatment to re-establish normal iron homeostasis for two weeks. All mice were then challenged with a lethal dose (1 X 10^3^) of the Type I CPS parental strain RH i.p and monitored for survival. As shown in Figure 6E, 5 X 10^4^ tachyzoites were not protective and 1 X 10^5^ were partially protective against lethal challenge. 5 X 10^5^ CPS tachyzoites induced near-complete protective immunity to lethal challenge. DFP induced iron restriction decreased CPS vaccine dose efficacy in both 1 X 10^4^ and 5 X 10^5^ CPS animals. This result demonstrates that iron is critical during immunization and iron restriction prevents development of immune memory to *T. gondii*.

### CD4+ T cell intrinsic CD71 is required for development of CD4+ T cell responses during T. gondii infection

Our results demonstrate that iron is critical for CD4+ and CD8+ T cell responses during *T. gondii* infection and this could be why DFP induced iron restriction results in higher morbidity, mortality and cyst burdens in the CNS. However, iron restriction appeared to impact CD4+ T cells more than CD8+ T cells. In addition, CD4+ T cells also upregulated CD71 more than CD8+ T cells after infection, and was further increased with DFP induced iron restriction. Therefore, we hypothesized that CD4+ T cell intrinsic CD71 was required for their activation and function. To test this hypothesis, we generated CD4- CreER^T2^ :: CD71^Flx/Flx^ mice for tamoxifen induced conditional knockout of CD71 in CD4+ T cells (Figure S4). CD4+ conditional CD71 knockout mice (CD4cCD71KO) were infected with ME49 cysts and the CD4+ and CD8+ T cell frequency and absolute numbers were measured by flow cytometry. The frequency and number of total CD4+ T cells in CD4cCD71KO compared to WT CreER^T2^ negative littermate controls was significantly reduced (Figure 7A, C). Surprisingly, the total CD8+ T cell population, not non-T cell populations, increased in frequency and absolute number in the CD4cCD71KO mice (Figure 7A, B, D). As further validation of the model, CD71 expression was significantly reduced on the CD4+ T cells in the CD4cCD71KO compared to WT littermate controls (Figure 7E). There was no change in TfR1 expression in CD8+ T cells (Figure 7F). To determine why the CD4+ T cell population was reduced we measured their proliferation using the proliferation marker Ki67 (67). Ki67 expression was reduced in CD4+ T cells from the CD4cCD71KO group by 52.7% compared to WT littermate controls (Figure 7G).

**Figure 7.**
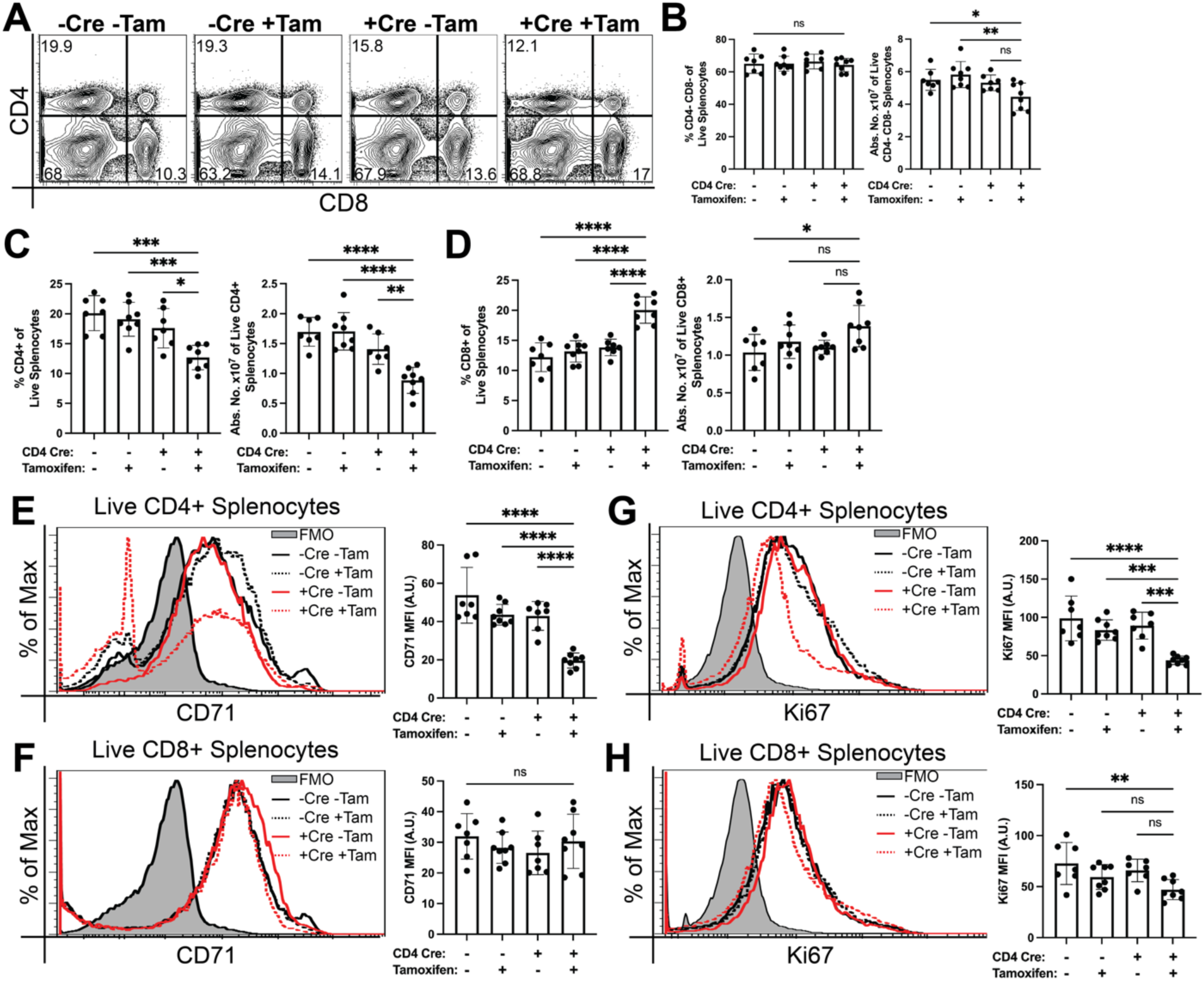
CD71 is required for activation and expansion of CD4+ T cells in a cell intrinsic manner during *T. gondii* infection. CD4cCD71KO mice and littermate controls were genotyped for possession (Cre+) or not (Cre-) of CreER^T2^ recombinase under the CD4 promotor, and all possessed flanking lox sites of the endogenous CD71 gene. Cre+ and Cre- littermates were alternatively treated (Tam+) or not (Tam-) with 100mg/kg tamoxifen i.p. daily for five days prior to oral infection with 8 cysts ME49. Splenocytes were harvested 10DPI for direct *ex vivo* staining according to the MIATA. **A-D)** Direct *ex vivo* staining of CD4 and CD8 is represented by contour plots and bar graphs presenting frequency and absolute numbers of B) non-T Cells (Live/CD4-/CD8-), C) CD4+ T cells (Live/CD4+/CD8-), and D) CD8+ T cells (Live/CD4-/CD8+). **E-F)** Direct *ex vivo* staining of CD71 is represented by histograms and bar graphs presenting geometric mean of CD71 intensity in E) CD4+ T Cells (Live/CD4+/CD8-) and F) CD8+ T cells (Live/CD4-/CD8+). **G-H)** Direct *ex vivo* staining of Ki67 is represented by histograms and bar graphs presenting geometric mean of Ki67 intensity in G) CD4+ T Cells (Live/CD4+/CD8-) and H) CD8+ T cells (Live/CD4-/CD8+). All data pooled from two independent experiments of N=3-4 per group. Bar graphs depict mean ± SD. Significance is denoted with * where p ≤ 0.05, ** where p ≤ 0.01, *** where p ≤ 0.005, **** where p ≤ 0.001, and NS where p > 0.05.

To examine how iron uptake via CD71 in CD4+ T cells impacted the immune response to *T. gondii*, we measured antigen experience (CD11aHiCD49d+) and functionality of CD4+ and CD8+ T cell subsets in CD4cCD71KO mice and WT littermate controls after ME49 infection. The frequency of antigen experienced CD4+ T cells in the CD4cCD71KO + tamoxifen group was 38.90%, significantly lower than the non-tamoxifen treated CD4cCD71KO (57.20%) and WT littermate controls with or without tamoxifen (53.56% and 58.01%, respectively) (Figure 8A). CD4+ T cells are required to maintain CD8+ T cell responses and memory development (26). To our surprise, the frequency and absolute number of antigen experienced CD11aHi CD49d+ CD8+ T Cells were not significantly different comparing CD4cCD71KO (58.33% in untreated and 64.84% in tamoxifien-treated) WT littermate controls (59.51% in untreated and 55.23% in tamoxifen-treated) (Figure 8B). The frequency and absolute number of polyfunctional CD4+ T cells was highly dependent on CD71 (Figure 8C). The polyfunctional (IFN*γ*+/TNF*α*+) CD4+ T cell proportions were reduced to 1.13% in tamoxifen-treated CD4cCD71KO from 11.46% and 6.78% in the littermate controls and 10.76% non-tamoxifen treated CD4cCD71KO (Figure 8C). There was no significant difference in the functional CD8+ T cell response across all groups (Figure 8D) suggesting that iron largely regulates the ability of the CD4+ T cells to expand and function during infection independent of CD4+ T cell help to developing CD8+ T cell responses. To further test this possibility we infected all mouse groups with ME49 and measured survival. No significant difference in survival was observed comparing CD4cCD71KO to WT litter mate controls with or without tamoxifen treatment (Figure 8E), supporting the importance of iron uptake for CD4 function, but not for helping CD8+ T cell responses develop.

**Figure 8.**
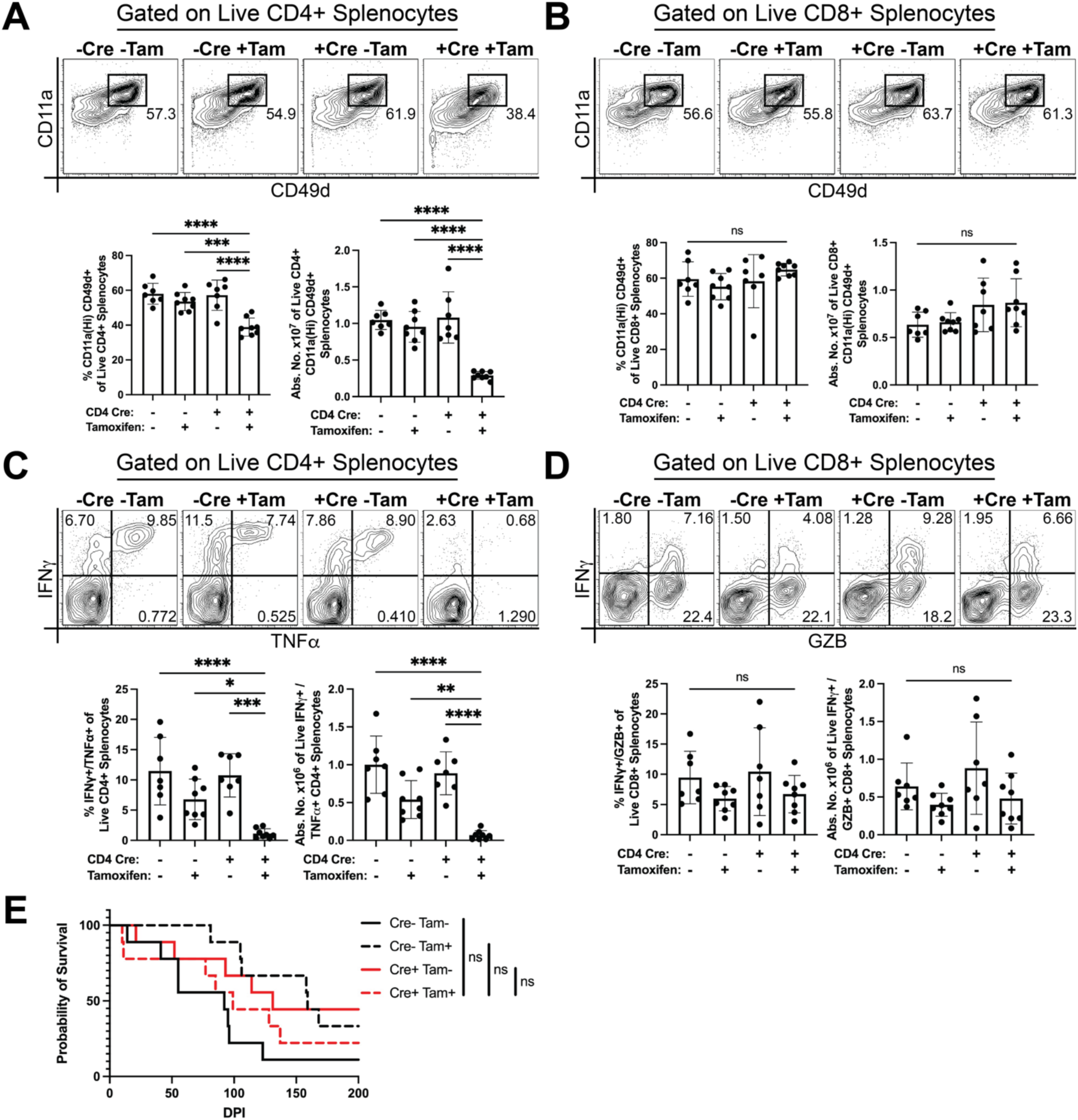
CD71 is required to increase antigen experience and polyfunctional responses of CD4+ T cells in *T. gondii* infection. CD4cCD71KO mice and littermate controls were genotyped for possession (Cre+) or not (Cre-) of CreER^T2^ recombinase under the CD4 promotor, and all possessed flanking lox sites of the endogenous CD71 gene. Cre+ and Cre- littermates were alternatively treated (Tam+) or not (Tam-) with 100mg/kg tamoxifen i.p. daily for five days prior to oral infection with 8 cysts ME49. Splenocytes were harvested 10DPI for stimulation or direct *ex vivo* staining according to the MIATA. **A)** Direct *ex vivo* staining of CD11a and CD49d is represented by contour plots and bar graphs presenting frequency and absolute numbers of antigen experienced CD4+ T cells (Live/CD4+/CD8-/CD11aHi/CD49d+). **B)** Direct *ex vivo* staining of CD11a and CD49d is represented by contour plots and bar graphs presenting frequency and absolute numbers of antigen experienced CD8+ T cells (Live/CD4-/CD8+/CD11aHi/CD49d). **C)** TLA-stimulated staining of IFN*γ* and TNF*α* is represented by contour plots and bar graphs presenting frequency and absolute numbers of polyfunctional CD4+ T Cells (Live/CD4+/CD8-/IFN*γ*+/TNF*α*+). **D)** TLA-stimulated staining of IFN*γ* is represented by contour plots and bar graphs presenting frequency and absolute numbers of polyfunctional CD8+ T Cells (Live/CD4-/CD8+/IFN*γ*+/GZB+). **E)** Survival of ME49 infected CD4cCD71KO mice and littermate controls treated with tamoxifen. Survival data pooled from two independent experiments of N=4-5 per group and analyzed by Kaplan-Meier statistics where NS indicates p > 0.05. All bar graphs pooled from two independent experiments of N=3-4 per group and depict mean ± SD. Bar graph significance is indicated with * where p ≤ 0.05, ** where p ≤ 0.01, *** where p ≤ 0.005, **** where p ≤ 0.001, and NS where p > 0.05.

## Discussion

Competition between host and pathogen for nutritional iron shapes the pathogenesis and outcome of infections. Here, we investigated this complex interaction during *T. gondii* infection. Our results demonstrate that iron is not only essential for *Toxoplasma gondii* replication, but also for optimal host immune responses because restricting iron *in vivo* results in higher morbidity and mortality in mice. During iron restriction with DFP, splenic and systemic IL-12 and IFN*γ* levels were significantly reduced in acute infection. However, in the absence of adaptive immune cells in RAG1-/- mice, iron restriction does not further enhance mortality due to infection. The immune cells impacted the most by iron restriction were CD4+ and CD8+ T cells, as antigen experience of both and CD4+ functionality were dependent on iron. Importantly, iron restriction inhibition of CD4+ and CD8+ T cells was independent of antigen load and was essential for development of vaccine induced long-term protection against lethal *T. gondii* challenge. Both CD4+ and CD8+ T cells upregulate CD71 protein expression in response to *T. gondii*, and especially so by antigen experienced CD4+ T cells when iron is limited. Lastly, we find that CD71 expression is required for expansion of antigen experienced and polyfunctional CD4+ T cell in a cell intrinsic manner. Taken together, these results suggest that iron homeostasis is a required nutrient for efficient CD4+ and CD8+ T cell responses to *T.* gondii and regulated by CD71 in CD4+ T cells. The implications of this study suggest that iron nutrition is an extenuating factor not only for resisting primary infections, but with long lasting effects even after iron balance is restored for vaccine efficacy.

The absolute requirement for iron in *T. gondii* biology remains an attractive target for therapeutic intervention, however, restricting iron could also have a negative effect on the development of immunity to this infection. The levels of IL-12 and IFN*γ* at 5DPI were reduced by iron restriction in both serum and spleen, but not completely absent. IL-12 is produced by the myeloid cell populations including neutrophils, monocytes, macrophages and dendritic cells during *T. gondii* infection (45, 46). Iron overload can affect the polarization and differentiation of monocytes into macrophages via mitochondrial oxidative stress (32, 33). Treatment of iron deficiency with intravenous iron can affect the differentiation of monocytes into macrophages and DCs (34). Although iron homeostasis could be important in these roles, we did not observe any significant difference in the ratios of these cells in the spleen during acute infection, suggesting that DFP induced iron restriction may not significantly impair myeloid cell cellular composition during *T. gondii* infection. IL-12-dependent IFN*γ* produced by NK cells is critical for early control of *T. gondii* and reaches its peak by day 5 post infection (22). Our results reveal NK cell responses in DFP induced iron restriction conditions remained intact at that time point compared to normal iron conditions. These results further support that iron may not have an important role in innate immunity to *T. gondii*. Taken together, this could indicate that parasite detection by the innate immune system was impaired and translates to lower cytokine production *in vivo,* but the potential capacity of NK cell IFN*γ* production is independent of iron homeostasis. Further exploration of the impact of iron in innate immunity to *T. gondii* is warranted to answer these questions.

Available iron and CD4+ T cell intrinsic CD71 are critical factors for the development of adaptive immunity to *T. gondii* infection. Blockade of CD71 transporting iron on human CD4+ T cells prevents their proliferation after anti-CD3/CD28 activation (38). Iron status can also affect the differentiation of T helper cell type, potentially impairing development of proper T helper populations important for controlling disease (37, 39, 40). Our results demonstrate that iron levels do impact the development of T cell responses during *T. gondii* infection. Expansion of antigen experienced CD11a(Hi) CD49d+ CD4+ and CD8+ T cells and CD4+ T cell functionality were inhibited in iron restriction conditions during *T. gondii* infection and after vaccination. CD4+ T cell intrinsic CD71 was essential for antigen experienced cell expansion and optimal polyfunctional responses. There are several possibilities that may explain the role of iron and CD71 in T cell expansion and activation during *T. gondii* infection.

Iron levels in a T cell can be sensed and could provide post transcriptional regulation of critical genes for T cell activation, proliferation, function, and differentiation. Iron is a known cofactor for Iron-Responsive Element (IRE) Binding Protein 1 (IRP1), which is a regulator of translation for many proteins, such as CD71, by differential binding of mRNA IRE’s (68). T cell transcription factors such as Tbx21, Eomesoderm, PRDM1, Akt/mTor, NF*κ*B, TCF1, and STATs, all critical for T cell activation, could contain IREs in their 5’ and 3’ UTRs that IRP1 could then bind to regulate their expression depending on cellular iron levels. There is evidence that at least Tbx21 and STAT1 may be mitigated by iron levels (69, 70).

T cell activation requires a metabolic shift to support their activation, proliferation, differentiation, and function to control diseases. Naïve, memory, and effector T cells exhibit distinct metabolic profiles (71–75) and undergoing a metabolic shift once they receive proper activation signals for antigen recognition, costimulation, and cytokines (76, 77). Iron and uptake via CD71 may supply sufficient iron levels for energetic demands of T cell activation. In addition to regulating mRNA stability and translation, IRP1 also works to regulate the citric acid cycle (TCA cycle) as aconitase when iron sulfur clusters are bound (78). Our study may have revealed how iron uptake is a critical fourth signal to turn on metabolic activity for CD4+ and CD8+ T cell expansion and function and how this is also critical for vaccine efficacy against this infection. Interestingly, our results also demonstrate that iron and CD71 mediated iron uptake is more important for CD4+ T cells to become activated than CD8+ T cells.

The activation, proliferation, differentiation and function of T cells can be regulated by critical second messengers: reactive oxygen species (ROS) produced inside of T cells. Iron and CD71 dependent iron uptake could impact ROS levels thereby affecting their intracellular signaling for activation. Iron reacts with hydrogen peroxide to produce hydroxy radicals (OH) by the Fenton reaction (79). The level of ROS and hydroxy radicals can be sensed by glutathione which then in turn regulates T cell proliferation (80). ROS can also be detected by the Nrf2-KEAP1-Cul3 which releases Nrf2 from the complex to become an active suppressor of T cell activation (80). Maintenance of moderate levels of ROS is also essential for proper TCR signaling that then can enhance T cell activation, proliferation, differentiation, and function (81–84). Understanding how iron and CD71 uptake impacts ROS signaling in T cells during *T. gondii* infection will be important future research.

CD71 may function independently of iron uptake in T cell activation. CD71 has been observed to co-localize with the T Cell Receptor (TCR) complex and to be a functional signaling component of the immune synapse (85). The mechanism(s) of how CD71 interacting with the TCR complex influence the level of T cell activation are unknown. CD71 is phosphorylated by Protein Kinase C (PKC) at position serine 24 in the cytoplasmic domain of CD71 away from the transferrin binding site (86, 87). Endocytosis of CD71 is required for iron uptake, but the serine 24 residue is dispensable for it, suggesting a potential still-undiscovered function of CD71 phosphorylation (88–90). Since PKC is central to TCR engagement downstream of CD3 and CD28 signaling (91), phosphorylation of CD71 by PKC would spatiotemporally position CD71 to contribute to signaling as part of the TCR complex. CD71 expression is ultimately controlled by IRP1-dependent iron sensing, therefore the secondary potential signaling function of CD71 in the TCR could act as another checkpoint for adequate iron levels to support T cell activation.

CD4+ and CD8+ T cells’ reliance on iron was different depending on strain of *T. gondii* infection. Our results indicate that iron may be more essential for regulating CD4+ T cell effector responses than CD8+ T cell effector responses during Type II *T. gondii* strain acute infection, but that effector responses of both CD4+ and CD8+ T cells are equally reliant on available iron after vaccination with the attenuated vaccine strain CPS. Antigen processing *T. gondii* infection can occur through cross presentation, phagocytosis, or autophagy (92–94). In typical type II ME49 infections, proliferation of the parasite is coupled with antigen processing, but in DFP induced iron limiting conditions, the parasite proliferation and dissemination are altered. This may differentially affect modes of antigen processing and presentation to CD4+ T cells compared to CD8+ T cells. For example, since iron levels influence autophagic potential of cells (95) and autophagy is a major route for MHCII antigen processing (96), it is plausible that modulation of autophagy by iron restriction preferentially inhibits antigen processing important for antigen experience for both CD4+ and CD8+ T cells but inhibits function of only CD4+ T cells in ME49 studies. In contrast, in CPS vaccination studies, the parasite cannot proliferate, thus, the potential differing effects of iron restriction on antigen processing and presentation to CD4+ or CD8+ T cells would be equal, providing some explanation as to why we observed inhibition of both CD4+ and CD8+ T cells in terms of antigen experience and function. Additionally, in both CPS and ME49 strain infections, the proportion of CD8+ FIII and FIV effector subpopulations, not the FII memory precursor subpopulation, were inhibited by iron restriction, suggesting that the type and strength of the CD8+ T cell response relies on iron as much as CD4+ T cell polyfunctional response.

In addition, CD4+ T cells may be more reliant on iron uptake by CD71 than CD8+ T cells. Our data may support this possibility because we observed that only antigen experienced CD4+ T cells upregulated their CD71 to a higher level in iron limiting conditions with DFP. CD4+ T cells are required to provide help for the activation, maintenance of activation, and memory cell differentiation of CD8+ T cells during *T. gondii* infection (24, 26, 66). The conditional knockout of CD71 in CD4+ T cells reduced overall frequencies and number of CD4+ T cells, CD4+ T cell antigen experience, and CD4+ functionality. These results demonstrate that CD71 is critical for development of CD4+ T cell responses to *T. gondii*. Based on these results we were expecting that there would be a decrease in CD8+ T cell responses due to the lack of CD4+ T cell help in the absence of CD71. However, this was not the case; within the CD8+ T cell compartment, there was no change of antigen experience or functionality in the CD4cCD71KO mouse treated with tamoxifen compared to all other groups. Intriguingly, the reduced frequency of total CD4+ T cells in the spleen in the CD4cCD71KO mice treated with tamoxifen was compensated by an increase in total CD8+ T cell frequencies and numbers. It is possible that the compensatory increase of overall CD8+ numbers could additionally compensate for reduced protection provided by CD4+ T cells in the conditional knockout, resulting in similar disease outcomes. A recent study has demonstrated that CD71 antibody blockade that inhibits iron uptake by human CD4+ T cells still results in generation of accessory helper T cells *in vitro* (38). An alternative explanation for our results then is that by knocking down expression of CD71 in CD4+ T cells, the generation of these accessory CD4+ T cell helper cells is retained. While overall functionality and expansion of antigen experienced CD4+ T cells is reduced, these cells might provide CD8+ T cells with the help they need to continue to function and develop memory. This is further supported by our survival data that shows no difference in long-term survival between CD4cCD71KO tamoxifen treated and WT litter mate controls. Further studies of the role of CD71 in the T cell responses to *T. gondii*, whether iron-dependent or -independent, will be helpful to uncover additional roles for iron as a fourth signal regulating T cell immunity during this infection.

In summary, the results of this study confirm the nutritional requirement of iron for both *Toxoplasma gondii* and its corresponding immune response and defined the adverse outcome of mice that do not meet this nutritional need. The mechanism driving the adverse outcome is explained by defective CD4+ and CD8+ T cell responses. Optimal CD4+ T cells responses were mediated by iron uptake receptor Transferrin Receptor 1. Many questions remain and will be addressed in future studies.

## List of Abbreviations

CD71: Transferrin Receptor 1
DFP: Deferiprone
TLA: Toxoplasma Lysate Antigen
(D,W)PI: (Days, Weeks) Post Infection
TCR: T Cell Receptor
MFI: Mean Fluorescent Intensity
ROS: Radical Oxygen Species
CD4cCD71KO: Conditional Knockout of CD71 in CD4+ T Cells

## Acknowledgements

The principal investigator and corresponding author of this project is J.P.G. Project conception was conducted by S.L.D. and J.P.G. Experiments were designed by S.L.D. and performed by S.L.D., T.R., H.K.K., and S.K.N. Data analysis was conducted by S.L.D., and T.R. Intellectual contributions were made by S.L.D. and J.P.G. Technical expertise was provided by J.P.G. Manuscript preparation was performed by S.L.D. and J.P.G., with contributions by all authors on revisions. Funding securement and project management was performed by J.P.G.

---

**Supplemental Figure 1.**
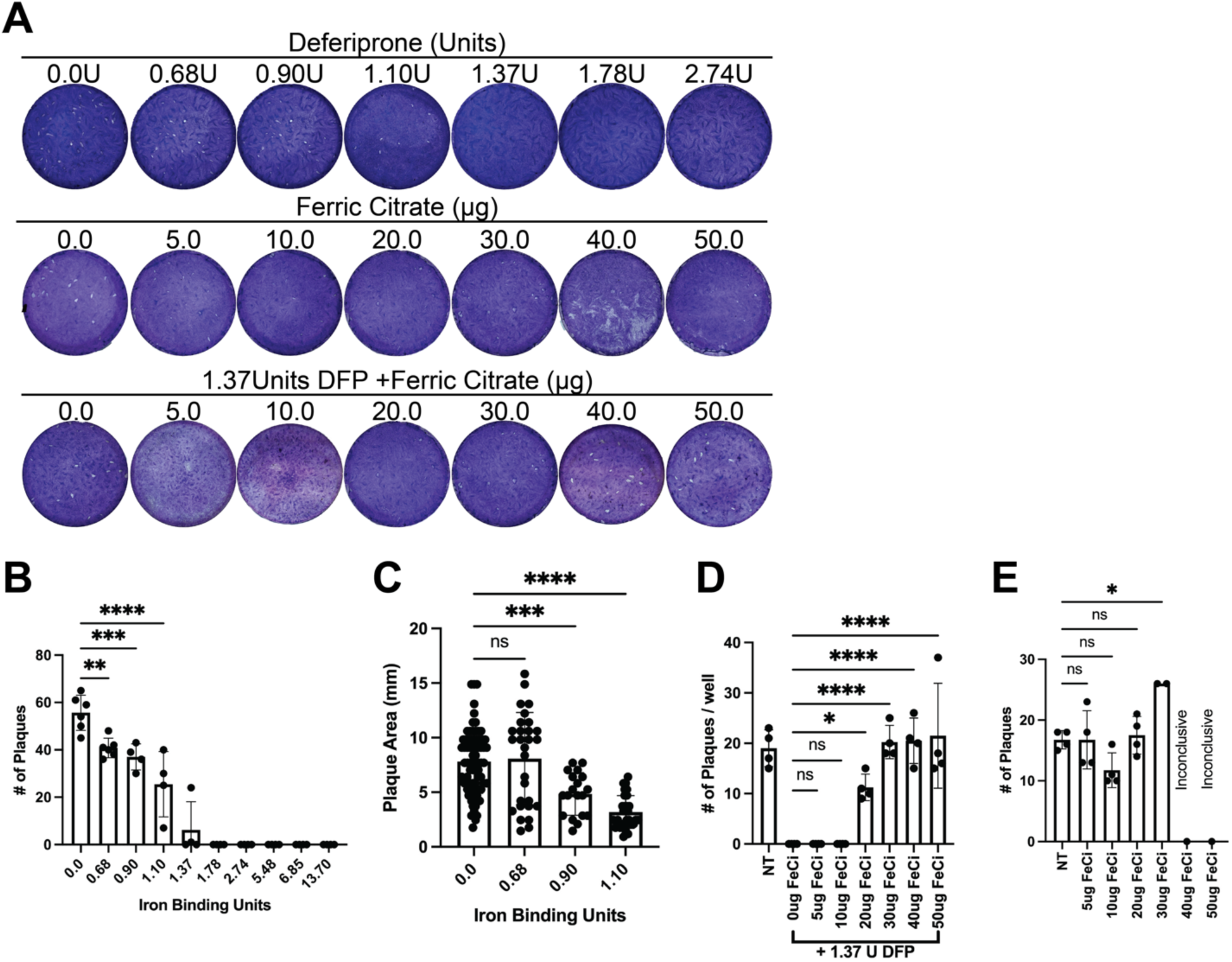
Titration of deferiprone and ferric citrate in plaque assays. The growth rate of RH strain *T. gondii* treated with indicated amounts of DFP or ferric citrate was determined by plaque assay. **A)** Shown are representative plaques formed by RH at indicated levels of DFP *(top),* ferric citrate *(middle)*, and ferric citrate + 1.37U DFP *(bottom).* **B)** Bar graph presents the number of RH plaques formed when treated with indicated doses of DFP. **C)** Bar graph presents the size of RH plaques formed when treated with indicated doses of DFP. **D)** Bar graph shows the number of plaques formed after supplementation of 1.37U DFP with indicated amounts of ferric citrate. **E)** Bar graph presents the number of plaques formed when supplemented with indicated amounts of ferric citrate alone. Data representative of two independent experiments. Bar graphs show mean ± SD. Significance is denoted with * where p ≤ 0.05, ** where p ≤ 0.01, *** where p ≤ 0.005, **** where p ≤ 0.001, and NS where p > 0.05.

**Supplemental Figure 2.**
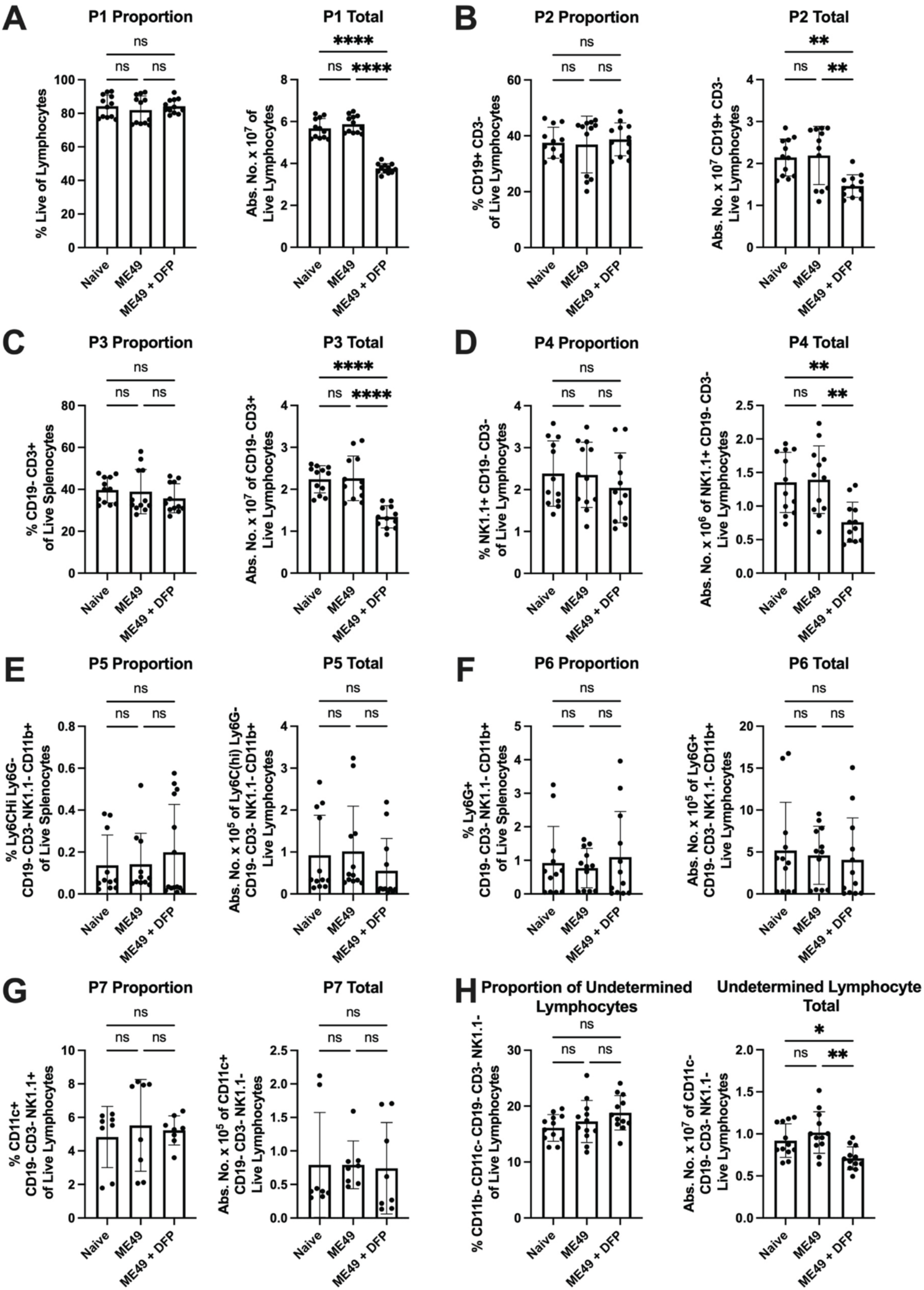
Iron restriction does not impact splenic proportions at 5DPI of *T. gondii* infection. C57BL/6 mice were infected orally with 8 cysts ME49 and treated with 75U DFP i.p. on days -1, 1, and 3. Splenocytes were isolated on day 5 for flow cytometric analysis. Bar graphs present frequency and absolute numbers of the following populations gated according to Figure 3A: **A)** P1 (Live Cells), **B)** P2 (Live, CD19+), **C)** P3 (Live, CD19+, CD3+, NK1.1-), **D)** P4 (Live, CD19-, CD3-, NK1.1+), **E)** P5 (Lineage-, CD11b+, Ly6CHi, Ly6G-, confirmed by side scatter profile), **F)** P6 (Lineage-, CD11b+ Ly6Cint, Ly6G+, confirmed by side scatter profile), **G)** P7 (Lineage-, CD11c+, I-A/I-E+), and **H)** P8 (Lineage- CD11b- CD11c-). Data pooled from three independent experiments. Bar graphs depict mean ± SD. Significance is denoted with * where p ≤ 0.05, ** where p ≤ 0.01, *** where p ≤ 0.005, **** where p ≤ 0.001, and NS where p > 0.05.

**Supplemental Figure 3.**
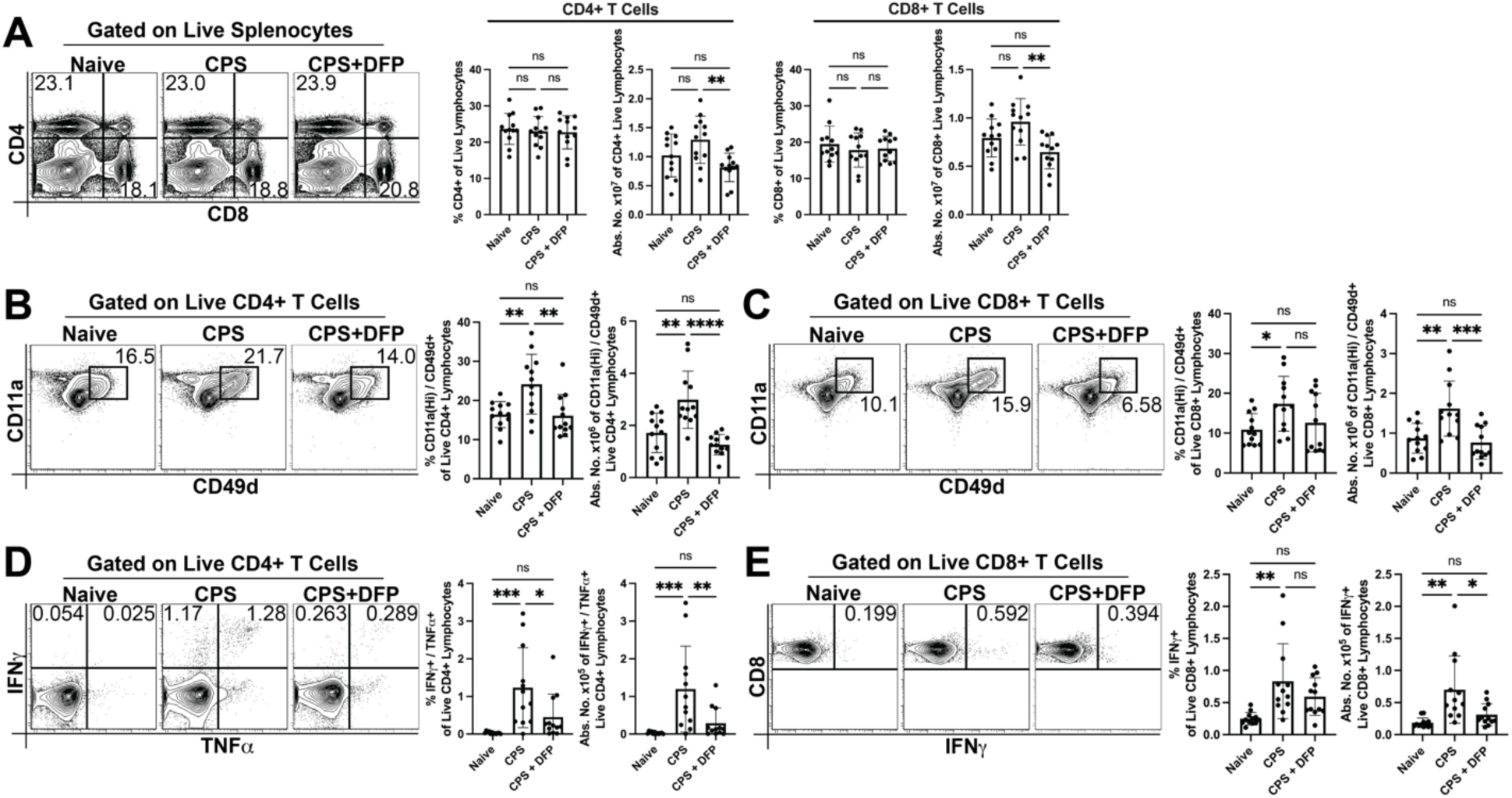
Inhibition of splenic T cell antigen experience and functionality by iron restriction during vaccination is independent of antigen load. C57BL/6 mice infected with 1 X 10^6^ tachyzoites CPS i.p. and treated with 75U DFP as described in the Materials and Methods. Animals were sacrificed and splenocytes were prepared for stimulation and for direct *ex vivo* staining. **A)** Direct *ex vivo* staining of CD4 and CD8 is represented by contour plots and bar graphs presenting frequency and absolute numbers of CD4+ T cells (Live/CD4+/CD8-) or CD8+ T Cells (Live/CD4-/CD8+). **B)** Direct *ex vivo* staining of CD11a and CD49d is represented by contour plots and bar graphs presenting frequency and absolute numbers of antigen experienced CD4+ T cells (Live/CD4+/CD8-/CD11aHi/CD49d+). **C)** Direct *ex vivo* staining of CD11a and CD49d is represented by contour plots and bar graphs presenting frequency and absolute numbers of antigen experienced CD8+ T cells (Live/CD4-/CD8+/CD11aHi/CD49d+). **D)** TLA- stimulated staining of IFN*γ* and TNF*α* is represented by contour plots and bar graphs of frequency and absolute numbers of polyfunctional CD4+ T Cells (Live/CD4+/CD8- /IFN*γ*+/TNF*α*+). **E)** TLA-stimulated staining of IFN*γ* is represented by contour plots and bar graphs of frequency and absolute numbers of functional CD8+ T Cells (Live/CD4- /CD8+/IFN*γ*+). All data pooled from three independent experiments of N=4 per group. Bar graphs depict mean ± SD. Significance is denoted with * where p ≤ 0.05, ** where p ≤ 0.01, *** where p ≤ 0.005, **** where p ≤ 0.001, and NS where p > 0.05.

**Supplemental Figure 4.**
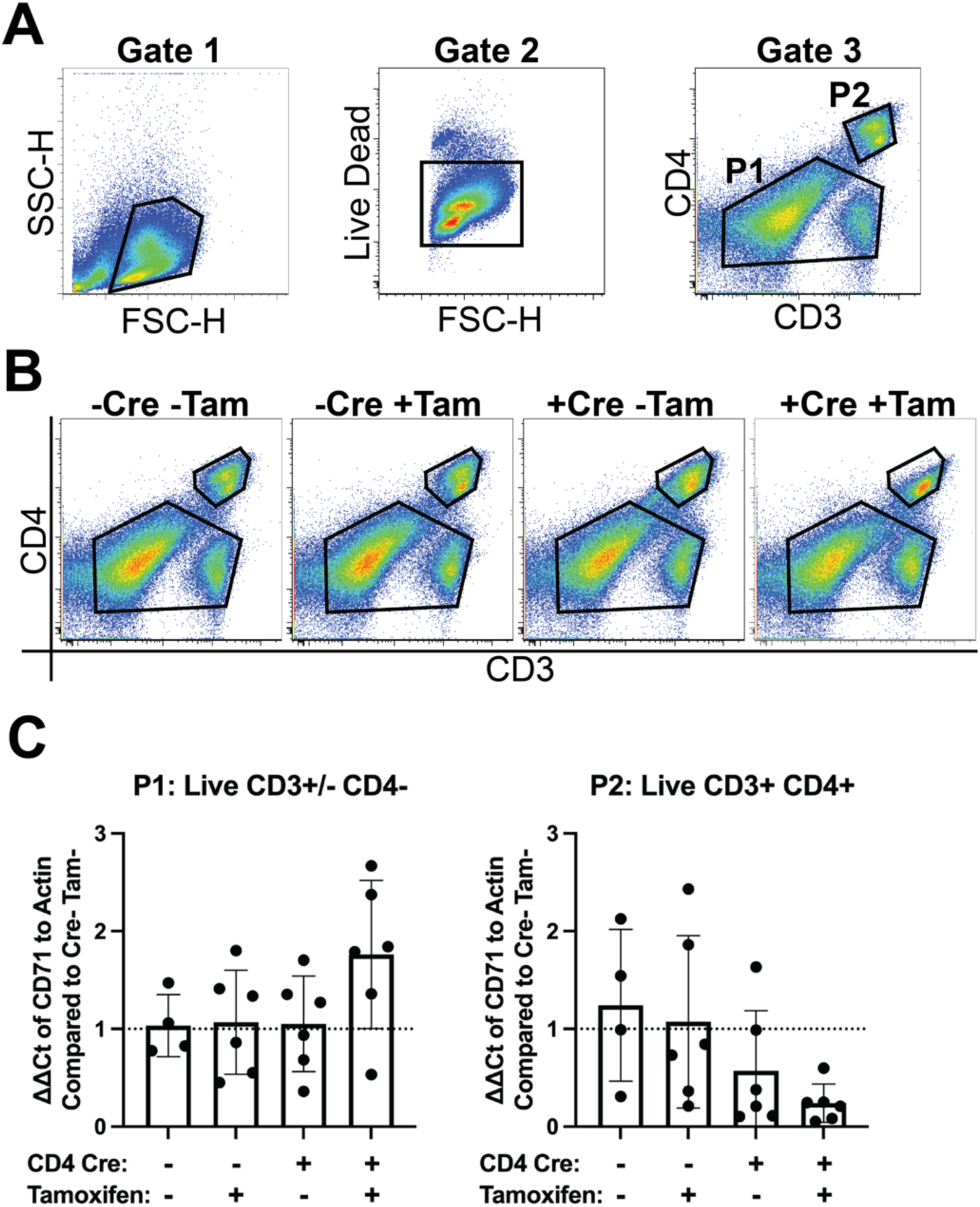
Conditional knockout of iron uptake receptor CD71 in CD4+ T cells. CD4cCD71KO mice and littermate controls were genotyped for possession (Cre+) or not (Cre-) of CreER^T2^ recombinase under the CD4 promotor, and all possessed flanking lox sites of the endogenous CD71 gene. Cre+ and Cre- littermates were alternatively treated (Tam+) or not (Tam-) with 100mg/kg tamoxifen i.p. daily for five days prior to oral infection with 8 cysts ME49. Splenocytes were harvested 10DPI for FACS and RT- PCR according to the MIATA. **A)** FACS gating strategy used for sorting of non CD4+ T cells (P1: Live, CD3+/-, CD4-) and CD4+ T cells (P2: Live, CD3+, CD4+). **B)** Pseudocolored dotplots represent FACS sorting populations for each group. **C)** Bar graphs present relative mRNA expression of CD71 compared to actin and normalized to the Cre- Tam- littermate control. The left graph shows CD71 mRNA expression of P1 sorted cells and the right graph shows CD71 mRNA expression of P2 sorted cells. All data was pooled from two independent experiments of N=1-4 per group and graphs are mean ± SD. No comparisons were statistically significant.

## References

1. Pappas, G., N. Roussos, and M. E. Falagas. 2009. Toxoplasmosis snapshots: global status of Toxoplasma gondii seroprevalence and implications for pregnancy and congenital toxoplasmosis. International journal for parasitology 39: 1385–1394.

2. Luft, B. J., and J. S. Remington. 1988. AIDS commentary. Toxoplasmic encephalitis. The Journal of infectious diseases 157: 1–6.

3. Tenter, A. M., A. R. Heckeroth, and L. M. Weiss. 2000. Toxoplasma gondii: from animals to humans. International journal for parasitology 30: 1217–1258.

4. Fox, B. A., J. P. Gigley, and D. J. Bzik. 2004. Toxoplasma gondii lacks the enzymes required for de novo arginine biosynthesis and arginine starvation triggers cyst formation. International journal for parasitology 34: 323–331.

5. Krug, E. C., J. J. Marr, and R. L. Berens. 1989. Purine metabolism in Toxoplasma gondii. J Biol Chem 264: 10601–10607.

6. Romano, J. D., S. Sonda, E. Bergbower, M. E. Smith, and I. Coppens. 2013. Toxoplasma gondii salvages sphingolipids from the host Golgi through the rerouting of selected Rab vesicles to the parasitophorous vacuole. Mol Biol Cell 24: 1974–1995.

7. Ma, E. H., G. Bantug, T. Griss, S. Condotta, R. M. Johnson, B. Samborska, N. Mainolfi, V. Suri, H. Guak, M. L. Balmer, M. J. Verway, T. C. Raissi, H. Tsui, G. Boukhaled, S. Henriques da Costa, C. Frezza, C. M. Krawczyk, A. Friedman, M. Manfredi, M. J. Richer, C. Hess, and R. G. Jones. 2017. Serine Is an Essential Metabolite for Effector T Cell Expansion. Cell metabolism 25: 345–357.

8. Tanaka, Y., Y. Nagai, M. Okumura, M. I. Greene, and T. Kambayashi. 2020. PRMT5 Is Required for T Cell Survival and Proliferation by Maintaining Cytokine Signaling. Front Immunol 11: 621.

9. Nagafuku, M., K. Okuyama, Y. Onimaru, A. Suzuki, Y. Odagiri, T. Yamashita, K. Iwasaki, M. Fujiwara, M. Takayanagi, I. Ohno, and J. Inokuchi. 2012. CD4 and CD8 T cells require different membrane gangliosides for activation. Proceedings of the National Academy of Sciences of the United States of America 109: E336–342.

10. Aghabi, D., M. Sloan, Z. Dou, A. J. Guerra, and C. R. Harding. 2021. The vacuolar iron transporter mediates iron detoxification in Toxoplasma gondii. bioRxiv.

11. Almeida, M. P. O., E. A. V. Ferro, M. P. P. Briceño, M. C. Oliveira, B. F. Barbosa, and N. M. Silva. 2019. Susceptibility of human villous (BeWo) and extravillous (HTR-8/SVneo) trophoblast cells to Toxoplasma gondii infection is modulated by intracellular iron availability. Parasitol Res 118: 1559–1572.

12. Dimier, I. H., and D. T. Bout. 1998. Interferon-gamma-activated primary enterocytes inhibit Toxoplasma gondii replication: a role for intracellular iron. Immunology 94: 488–495.

13. Byrd, T. F., and M. A. Horwitz. 1993. Regulation of transferrin receptor expression and ferritin content in human mononuclear phagocytes. Coordinate upregulation by iron transferrin and downregulation by interferon gamma. The Journal of clinical investigation 91: 969–976.

14. Gail, M., U. Gross, and W. Bohne. 2004. Transferrin receptor induction in Toxoplasma gondii-infected HFF is associated with increased iron-responsive protein 1 activity and is mediated by secreted factors. Parasitol Res 94: 233–239.

15. Oliveira, M. C., L. B. Coutinho, M. P. O. Almeida, M. P. Briceno, E. C. B. Araujo, and N. M. Silva. 2020. The Availability of Iron Is Involved in the Murine Experimental Toxoplasma gondii Infection Outcome. Microorganisms 8.

16. Plattner, F., F. Yarovinsky, S. Romero, D. Didry, M. F. Carlier, A. Sher, and D. Soldati-Favre. 2008. Toxoplasma profilin is essential for host cell invasion and TLR11-dependent induction of an interleukin-12 response. Cell host & microbe 3: 77–87.

17. Yarovinsky, F., S. Hieny, and A. Sher. 2008. Recognition of Toxoplasma gondii by TLR11 prevents parasite-induced immunopathology. Journal of immunology (Baltimore, Md. : 1950) 181: 8478–8484.

18. Gazzinelli, R. T., M. Wysocka, S. Hayashi, E. Y. Denkers, S. Hieny, P. Caspar, G. Trinchieri, and A. Sher. 1994. Parasite-induced IL-12 stimulates early IFN-gamma synthesis and resistance during acute infection with Toxoplasma gondii. Journal of immunology (Baltimore, Md. : 1950) 153: 2533–2543.

19. Chen, M., F. Aosai, K. Norose, H. S. Mun, O. Takeuchi, S. Akira, and A. Yano. 2002. Involvement of MyD88 in host defense and the down-regulation of anti-heat shock protein 70 autoantibody formation by MyD88 in Toxoplasma gondii-infected mice. The Journal of parasitology 88: 1017–1019.

20. Hitziger, N., I. Dellacasa, B. Albiger, and A. Barragan. 2005. Dissemination of Toxoplasma gondii to immunoprivileged organs and role of Toll/interleukin-1 receptor signalling for host resistance assessed by in vivo bioluminescence imaging. Cell Microbiol 7: 837–848.

21. Hunter, C. A., C. S. Subauste, V. H. Van Cleave, and J. S. Remington. 1994. Production of gamma interferon by natural killer cells from Toxoplasma gondii-infected SCID mice: regulation by interleukin-10, interleukin-12, and tumor necrosis factor alpha. Infection and immunity 62: 2818–2824.

22. Gazzinelli, R. T., S. Hieny, T. A. Wynn, S. Wolf, and A. Sher. 1993. Interleukin 12 is required for the T-lymphocyte-independent induction of interferon gamma by an intracellular parasite and induces resistance in T-cell-deficient hosts. Proceedings of the National Academy of Sciences of the United States of America 90: 6115–6119.

23. López-Yglesias, A. H., E. Burger, E. Camanzo, A. T. Martin, A. M. Araujo, S. F. Kwok, and F. Yarovinsky. 2021. T-bet-dependent ILC1- and NK cell-derived IFN-γ mediates cDC1-dependent host resistance against Toxoplasma gondii. PLoS pathogens 17: e1008299.

24. Gigley, J. P., B. A. Fox, and D. J. Bzik. 2009. Cell-mediated immunity to Toxoplasma gondii develops primarily by local Th1 host immune responses in the absence of parasite replication. Journal of immunology (Baltimore, Md. : 1950) 182: 1069–1078.

25. Gigley, J. P., B. A. Fox, and D. J. Bzik. 2009. Long-term immunity to lethal acute or chronic type II Toxoplasma gondii infection is effectively induced in genetically susceptible C57BL/6 mice by immunization with an attenuated type I vaccine strain. Infection and immunity 77: 5380–5388.

26. Casciotti, L., K. H. Ely, M. E. Williams, and I. A. Khan. 2002. CD8(+)-T-cell immunity against Toxoplasma gondii can be induced but not maintained in mice lacking conventional CD4(+) T cells. Infection and immunity 70: 434–443.

27. Bhadra, R., J. P. Gigley, L. M. Weiss, and I. A. Khan. 2011. Control of Toxoplasma reactivation by rescue of dysfunctional CD8+ T-cell response via PD-1-PDL-1 blockade. Proceedings of the National Academy of Sciences of the United States of America 108: 9196–9201.

28. Ivanova, D. L., R. Krempels, S. L. Denton, K. D. Fettel, G. M. Saltz, D. Rach, R. Fatima, T. Mundhenke, J. Materi, I. R. Dunay, and J. P. Gigley. 2020. NK Cells Negatively Regulate CD8 T Cells to Promote Immune Exhaustion and Chronic Toxoplasma gondii Infection. Frontiers in cellular and infection microbiology 10: 313.

29. Xiao, J., Y. Li, R. H. Yolken, and R. P. Viscidi. 2018. PD-1 immune checkpoint blockade promotes brain leukocyte infiltration and diminishes cyst burden in a mouse model of Toxoplasma infection. Journal of neuroimmunology 319: 55–62.

30. Hwang, S., D. A. Cobb, R. Bhadra, B. Youngblood, and I. A. Khan. 2016. Blimp-1-mediated CD4 T cell exhaustion causes CD8 T cell dysfunction during chronic toxoplasmosis. The Journal of experimental medicine 213: 1799–1818.

31. Kono, M., S. Matsuhiroya, A. Obuchi, T. Takahashi, S. Imoto, S. Kawano, and K. Saigo. 2020. Deferasirox, an iron-chelating agent, alleviates acute lung inflammation by inhibiting neutrophil activation and extracellular trap formation. J Int Med Res 48: 300060520951015.

32. Agoro, R., M. Taleb, V. F. J. Quesniaux, and C. Mura. 2018. Cell iron status influences macrophage polarization. PloS one 13: e0196921.

33. Cui, Y., S. Gutierrez, S. Ariai, L. Öberg, K. Thörn, U. Gehrmann, S. M. Cloonan, T. Naessens, and H. Olsson. 2022. Non-heme iron overload impairs monocyte to macrophage differentiation via mitochondrial oxidative stress. Frontiers in Immunology 13.

34. Fell, L. H., S. Seiler-Mussler, A. B. Sellier, B. Rotter, P. Winter, M. Sester, D. Fliser, G. H. Heine, and A. M. Zawada. 2016. Impact of individual intravenous iron preparations on the differentiation of monocytes towards macrophages and dendritic cells. Nephrol Dial Transplant 31: 1835–1845.

35. Flynn, M. G., L. Mackinnon, V. Gedge, M. Fahlman, and T. Brickman. 2003. Influence of iron status and iron supplements on natural killer cell activity in trained women runners. Int J Sports Med 24: 217–222.

36. Baines, M. G., F. L. Lafleur, and B. E. Holbein. 1983. Involvement of transferrin and transferrin receptors in human natural killer effector:target interaction. Immunol Lett 7: 51–55.

37. Yarosz, E. L., C. Ye, A. Kumar, C. Black, E. K. Choi, Y. A. Seo, and C. H. Chang. 2020. Cutting Edge: Activation-Induced Iron Flux Controls CD4 T Cell Proliferation by Promoting Proper IL-2R Signaling and Mitochondrial Function. Journal of immunology (Baltimore, Md. : 1950) 204: 1708–1713.

38. Berg, V., M. Modak, J. Brell, A. Puck, S. Kunig, S. Jutz, P. Steinberger, G. J. Zlabinger, and J. Stockl. 2020. Iron Deprivation in Human T Cells Induces Nonproliferating Accessory Helper Cells. Immunohorizons 4: 165–177.

39. Pfeifhofer-Obermair, C., P. Tymoszuk, M. Nairz, A. Schroll, G. Klais, E. Demetz, S. Engl, N. Brigo, and G. Weiss. 2021. Regulation of Th1 T Cell Differentiation by Iron via Upregulation of T Cell Immunoglobulin and Mucin Containing Protein-3 (TIM-3). Front Immunol 12: 637809.

40. Leung, S., A. Holbrook, B. King, H. T. Lu, V. Evans, N. Miyamoto, C. Mallari, S. Harvey, D. Davey, E. Ho, W. W. Li, J. Parkinson, R. Horuk, S. Jaroch, M. Berger, W. Skuballa, C. West, R. Pulk, G. Phillips, J. Bryant, B. Subramanyam, C. Schaefer, H. Salamon, E. Lyons, D. Schilling, H. Seidel, J. Kraetzschmar, M. Snider, and D. Perez. 2005. Differential inhibition of inducible T cell cytokine secretion by potent iron chelators. J Biomol Screen 10: 157–167.

41. Rodriguez, R., C. L. Jung, V. Gabayan, J. C. Deng, T. Ganz, E. Nemeth, and Y. Bulut. 2014. Hepcidin induction by pathogens and pathogen-derived molecules is strongly dependent on interleukin-6. Infection and immunity 82: 745–752.

42. Chen, A. C., A. Donovan, R. Ned-Sykes, and N. C. Andrews. 2015. Noncanonical role of transferrin receptor 1 is essential for intestinal homeostasis. Proceedings of the National Academy of Sciences of the United States of America 112: 11714–11719.

43. Hatcher, H. C., R. N. Singh, F. M. Torti, and S. V. Torti. 2009. Synthetic and natural iron chelators: therapeutic potential and clinical use. Future Med Chem 1: 1643–1670.

44. Adams, L. B., J. B. Hibbs, Jr., R. R. Taintor, and J. L. Krahenbuhl. 1990. Microbiostatic effect of murine-activated macrophages for Toxoplasma gondii. Role for synthesis of inorganic nitrogen oxides from L-arginine. Journal of immunology (Baltimore, Md. : 1950) 144: 2725–2729.

45. Liu, C.-H., Y.-t. Fan, A. Dias, L. Esper, R. A. Corn, A. Bafica, F. S. Machado, and J. Aliberti. 2006. Cutting Edge: Dendritic Cells Are Essential for In Vivo IL-12 Production and Development of Resistance against Toxoplasma gondii Infection in Mice. The Journal of Immunology 177: 31–35.

46. Mercer, H. L., L. M. Snyder, C. M. Doherty, B. A. Fox, D. J. Bzik, and E. Y. Denkers. 2020. Toxoplasma gondii dense granule protein GRA24 drives MyD88-independent p38 MAPK activation, IL-12 production and induction of protective immunity. PLoS pathogens 16: e1008572.

47. Mombaerts, P., J. Iacomini, R. S. Johnson, K. Herrup, S. Tonegawa, and V. E. Papaioannou. 1992. RAG-1-deficient mice have no mature B and T lymphocytes. Cell 68: 869–877.

48. Hunter, C. A., R. Chizzonite, and J. S. Remington. 1995. Il-1-Beta Is Required for Il-12 to Induce Production of Ifn-Gamma by Nk-Cells - a Role for Il-1-Beta in the T-Cell-Independent Mechanism of Resistance against Intracellular Pathogens. Journal of Immunology 155: 4347–4354.

49. Goldszmid, R. S., P. Caspar, A. Rivollier, S. White, A. Dzutsev, S. Hieny, B. Kelsall, G. Trinchieri, and A. Sher. 2012. NK cell-derived interferon-gamma orchestrates cellular dynamics and the differentiation of monocytes into dendritic cells at the site of infection. Immunity 36: 1047–1059.

50. Lee, Y. H., J. Y. Channon, T. Matsuura, J. D. Schwartzman, D. W. Shin, and L. H. Kasper. 1999. Functional and quantitative analysis of splenic T cell immune responses following oral toxoplasma gondii infection in mice. Experimental parasitology 91: 212–221.

51. Carrof, B., M. Levacher-Clergeot, F. Chau, J. J. Pocidalo, and F. Derouin. 1994. Toxoplasma gondii: kinetics of lymphocyte subsets in blood and spleen of perorally infected mice. Experimental parasitology 78: 410–417.

52. McDermott, D. S., and S. M. Varga. 2011. Quantifying antigen-specific CD4 T cells during a viral infection: CD4 T cell responses are larger than we think. Journal of immunology (Baltimore, Md. : 1950) 187: 5568–5576.

53. Haque, S., J. Franck, H. Dumon, L. H. Kasper, and A. Haque. 1999. Protection against lethal toxoplasmosis in mice by an avirulent strain of Toxoplasma gondii: stimulation of IFN-gamma and TNF-alpha response. Experimental parasitology 93: 231–240.

54. Gazzinelli, R. T., I. Eltoum, T. A. Wynn, and A. Sher. 1993. Acute cerebral toxoplasmosis is induced by in vivo neutralization of TNF-alpha and correlates with the down-regulated expression of inducible nitric oxide synthase and other markers of macrophage activation. Journal of immunology (Baltimore, Md. : 1950) 151: 3672–3681.

55. Shirahata, T., T. Yamashita, C. Ohta, H. Goto, and A. Nakane. 1994. CD8+ T lymphocytes are the major cell population involved in the early gamma interferon response and resistance to acute primary Toxoplasma gondii infection in mice. Microbiol Immunol 38: 789–796.

56. Jongert, E., A. Lemiere, J. Van Ginderachter, S. De Craeye, K. Huygen, and S. D’Souza. 2010. Functional characterization of in vivo effector CD4(+) and CD8(+) T cell responses in acute Toxoplasmosis: an interplay of IFN-gamma and cytolytic T cells. Vaccine 28: 2556–2564.

57. Denkers, E. Y., and R. T. Gazzinelli. 1998. Regulation and function of T-cell-mediated immunity during Toxoplasma gondii infection. Clinical microbiology reviews 11: 569–588.

58. Giannetti, A. M., P. J. Halbrooks, A. B. Mason, T. M. Vogt, C. A. Enns, and P. J. Bjorkman. 2005. The molecular mechanism for receptor-stimulated iron release from the plasma iron transport protein transferrin. Structure 13: 1613–1623.

59. Bayer, A. L., P. Baliga, and J. E. Woodward. 1998. Transferrin receptor in T cell activation and transplantation. Journal of leukocyte biology 64: 19–24.

60. Brekelmans, P., P. van Soest, P. J. Leenen, and W. van Ewijk. 1994. Inhibition of proliferation and differentiation during early T cell development by anti-transferrin receptor antibody. European journal of immunology 24: 2896–2902.

61. Brekelmans, P., P. van Soest, J. Voerman, P. P. Platenburg, P. J. Leenen, and W. van Ewijk. 1994. Transferrin receptor expression as a marker of immature cycling thymocytes in the mouse. Cellular immunology 159: 331–339.

62. Wilson, D. C., S. Matthews, and G. S. Yap. 2008. IL-12 signaling drives CD8+ T cell IFN-gamma production and differentiation of KLRG1+ effector subpopulations during Toxoplasma gondii Infection. Journal of immunology (Baltimore, Md. : 1950) 180: 5935–5945.

63. Sanecka, A., N. Yoshida, E. M. Kolawole, H. Patel, B. D. Evavold, and E. M. Frickel. 2018. T Cell Receptor-Major Histocompatibility Complex Interaction Strength Defines Trafficking and CD103(+) Memory Status of CD8 T Cells in the Brain. Front Immunol 9: 1290.

64. Moretto, M. M., J. Chen, M. Meador, J. Phan, and I. A. Khan. 2023. A Lower Dose of Infection Generates a Better Long-Term Immune Response against Toxoplasma gondii. Immunohorizons 7: 177–190.

65. Fox, B. A., and D. J. Bzik. 2002. De novo pyrimidine biosynthesis is required for virulence of Toxoplasma gondii. Nature 415: 926–929.

66. Jordan, K. A., E. H. Wilson, E. D. Tait, B. A. Fox, D. S. Roos, D. J. Bzik, F. Dzierszinski, and C. A. Hunter. 2009. Kinetics and phenotype of vaccine-induced CD8+ T-cell responses to Toxoplasma gondii. Infection and immunity 77: 3894–3901.

67. Gerdes, J., H. Lemke, H. Baisch, H. H. Wacker, U. Schwab, and H. Stein. 1984. Cell cycle analysis of a cell proliferation-associated human nuclear antigen defined by the monoclonal antibody Ki-67. Journal of immunology (Baltimore, Md. : 1950) 133: 1710–1715.

68. Hentze, M. W., and L. C. Kuhn. 1996. Molecular control of vertebrate iron metabolism: mRNA-based regulatory circuits operated by iron, nitric oxide, and oxidative stress. Proceedings of the National Academy of Sciences of the United States of America 93: 8175–8182.

69. Van Den Ham, K. M., M. T. Shio, A. Rainone, S. Fournier, C. M. Krawczyk, and M. Olivier. 2015. Iron prevents the development of experimental cerebral malaria by attenuating CXCR3-mediated T cell chemotaxis. PloS one 10: e0118451.

70. Regis, G., M. Bosticardo, L. Conti, S. De Angelis, D. Boselli, B. Tomaino, P. Bernabei, M. Giovarelli, and F. Novelli. 2005. Iron regulates T-lymphocyte sensitivity to the IFN-gamma/STAT1 signaling pathway in vitro and in vivo. Blood 105: 3214–3221.

71. Rathmell, J. C., M. G. Vander Heiden, M. H. Harris, K. A. Frauwirth, and C. B. Thompson. 2000. In the absence of extrinsic signals, nutrient utilization by lymphocytes is insufficient to maintain either cell size or viability. Mol Cell 6: 683–692.

72. Frauwirth, K. A., J. L. Riley, M. H. Harris, R. V. Parry, J. C. Rathmell, D. R. Plas, R. L. Elstrom, C. H. June, and C. B. Thompson. 2002. The CD28 signaling pathway regulates glucose metabolism. Immunity 16: 769–777.

73. Verbist, K. C., C. S. Guy, S. Milasta, S. Liedmann, M. M. Kaminski, R. Wang, and D. R. Green. 2016. Metabolic maintenance of cell asymmetry following division in activated T lymphocytes. Nature 532: 389–393.

74. Menk, A. V., N. E. Scharping, R. S. Moreci, X. Zeng, C. Guy, S. Salvatore, H. Bae, J. Xie, H. A. Young, S. G. Wendell, and G. M. Delgoffe. 2018. Early TCR Signaling Induces Rapid Aerobic Glycolysis Enabling Distinct Acute T Cell Effector Functions. Cell Rep 22: 1509–1521.

75. Loschinski, R., M. Bottcher, A. Stoll, H. Bruns, A. Mackensen, and D. Mougiakakos. 2018. IL-21 modulates memory and exhaustion phenotype of T-cells in a fatty acid oxidation-dependent manner. Oncotarget 9: 13125–13138.

76. Wang, R., Christopher P. Dillon, Lewis Z. Shi, S. Milasta, R. Carter, D. Finkelstein, Laura L. McCormick, P. Fitzgerald, H. Chi, J. Munger, and Douglas R. Green. 2011. The Transcription Factor Myc Controls Metabolic Reprogramming upon T Lymphocyte Activation. Immunity 35: 871–882.

77. Frauwirth, K. A., J. L. Riley, M. H. Harris, R. V. Parry, J. C. Rathmell, D. R. Plas, R. L. Elstrom, C. H. June, and C. B. Thompson. 2002. The CD28 Signaling Pathway Regulates Glucose Metabolism. Immunity 16: 769–777.

78. Gray, N. K., S. Quick, B. Goossen, A. Constable, H. Hirling, L. C. Kühn, and M. W. Hentze. 1993. Recombinant iron-regulatory factor functions as an iron-responsive-element-binding protein, a translational repressor and an aconitase. A functional assay for translational repression and direct demonstration of the iron switch. Eur J Biochem 218: 657–667.

79. Silva-Gomes, S., S. Vale-Costa, R. Appelberg, and M. S. Gomes. 2013. Iron in intracellular infection: to provide or to deprive? Frontiers in cellular and infection microbiology 3: 96.

80. Yarosz, E. L., and C. H. Chang. 2018. The Role of Reactive Oxygen Species in Regulating T Cell-mediated Immunity and Disease. Immune Netw 18: e14.

81. Sena, L. A., S. Li, A. Jairaman, M. Prakriya, T. Ezponda, D. A. Hildeman, C.-R. Wang, P. T. Schumacker, J. D. Licht, and H. Perlman. 2013. Mitochondria are required for antigen-specific T cell activation through reactive oxygen species signaling. Immunity 38: 225–236.

82. Lepez, A., T. Pirnay, S. Denanglaire, D. Perez-Morga, M. Vermeersch, O. Leo, and F. Andris. 2020. Long-term T cell fitness and proliferation is driven by AMPK-dependent regulation of reactive oxygen species. Sci Rep 10: 21673.

83. Kaminski, M. M., S. W. Sauer, C. D. Klemke, D. Süss, J. G. Okun, P. H. Krammer, and K. Gülow. 2010. Mitochondrial reactive oxygen species control T cell activation by regulating IL-2 and IL-4 expression: mechanism of ciprofloxacin-mediated immunosuppression. Journal of immunology (Baltimore, Md. : 1950) 184: 4827–4841.

84. Arbore, G., E. E. West, R. Spolski, A. A. B. Robertson, A. Klos, C. Rheinheimer, P. Dutow, T. M. Woodruff, Z. X. Yu, L. A. O’Neill, R. C. Coll, A. Sher, W. J. Leonard, J. Köhl, P. Monk, M. A. Cooper, M. Arno, B. Afzali, H. J. Lachmann, A. P. Cope, K. D. Mayer-Barber, and C. Kemper. 2016. T helper 1 immunity requires complement-driven NLRP3 inflammasome activity in CD4⁺ T cells. Science 352: aad1210.

85. Salmeron, A., A. Borroto, M. Fresno, M. J. Crumpton, S. C. Ley, and B. Alarcon.1995. Transferrin receptor induces tyrosine phosphorylation in T cells and is physically associated with the TCR zeta-chain. Journal of immunology (Baltimore, Md. : 1950) 154: 1675–1683.

86. Davis, R. J., G. L. Johnson, D. J. Kelleher, J. K. Anderson, J. E. Mole, and M. P. Czech. 1986. Identification of serine 24 as the unique site on the transferrin receptor phosphorylated by protein kinase C. J Biol Chem 261: 9034–9041.

87. Cheng, Y., O. Zak, P. Aisen, S. C. Harrison, and T. Walz. 2004. Structure of the human transferrin receptor-transferrin complex. Cell 116: 565–576.

88. Rothenberger, S., B. J. Iacopetta, and L. C. Kuhn. 1987. Endocytosis of the transferrin receptor requires the cytoplasmic domain but not its phosphorylation site. Cell 49: 423–431.

89. Zerial, M., M. Suomalainen, M. Zanetti-Schneider, C. Schneider, and H. Garoff. 1987. Phosphorylation of the human transferrin receptor by protein kinase C is not required for endocytosis and recycling in mouse 3T3 cells. The EMBO journal 6: 2661–2667.

90. Davis, R. J., and H. Meisner. 1987. Regulation of transferrin receptor cycling by protein kinase C is independent of receptor phosphorylation at serine 24 in Swiss 3T3 fibroblasts. J Biol Chem 262: 16041–16047.

91. Isakov, N., and A. Altman. 2002. Protein kinase C(theta) in T cell activation. Annu Rev Immunol 20: 761–794.

92. Goldszmid, R. S., I. Coppens, A. Lev, P. Caspar, I. Mellman, and A. Sher. 2009. Host ER-parasitophorous vacuole interaction provides a route of entry for antigen cross-presentation in Toxoplasma gondii-infected dendritic cells. The Journal of experimental medicine 206: 399–410.

93. Lee, Y., M. Sasai, J. S. Ma, N. Sakaguchi, J. Ohshima, H. Bando, T. Saitoh, S. Akira, and M. Yamamoto. 2015. p62 Plays a Specific Role in Interferon-γ-Induced Presentation of a Toxoplasma Vacuolar Antigen. Cell Rep 13: 223–233.

94. Yano, A., S. Ohno, K. Norose, T. Baba, K. Yamashita, F. Aosai, and K. Segawa. 1992. Antigen presentation by Toxoplasma-infected cells: antigen entry through cell membrane fusion. Int Arch Allergy Immunol 98: 13–17.

95. Wang, Y., M. Wang, Y. Liu, H. Tao, S. Banerjee, S. Srinivasan, E. Nemeth, M. J. Czaja, and P. He. 2022. Integrated regulation of stress responses, autophagy and survival by altered intracellular iron stores. Redox Biol 55: 102407.

96. Romao, S., N. Gasser, A. C. Becker, B. Guhl, M. Bajagic, D. Vanoaica, U. Ziegler, J. Roesler, J. Dengjel, and J. Reichenbach. 2013. Autophagy proteins stabilize pathogen-containing phagosomes for prolonged MHC II antigen processing. Journal of Cell Biology 203: 757–766.

